# Semantic Information Orthogonal to Visual Features Peaks in Lateral Occipitotemporal Cortex

**DOI:** 10.64898/2026.03.14.711805

**Authors:** Arun Ram Ponnambalam, Krishnan Pottore Venkiteswaran

## Abstract

Language model embeddings of scene descriptions predict responses in the human higher visual cortex. However, a fundamental question remains: does this alignment reflect truly visually-independent semantic content, or does it occur because language models better mimic the complex visual features that drive these areas? We used 7T fMRI data from the Natural Scenes Dataset to directly address this by removing the influence of visual feature embeddings from language model embeddings, isolating semantic content that is separate from the visual signal. We then used these visually-independent embeddings to predict brain responses in individual voxels through cross-validated ridge regression. After adjusting for visual signals, we found a clear difference in brain regions: the lateral occipitotemporal cortex, especially in areas selective for body perception, showed significantly more visually-independent semantic variance compared to ventral stream regions. In contrast, the early visual cortex displayed notably negative predictions after adjustment, confirming that our method effectively removed visually-driven signals. This pattern was consistent across all eight subjects, both hemispheres, and six combinations of language models and visual feature architectures. These findings suggest that the lateral stream retains substantially more variance from language models unrelated to various visual feature models than the ventral stream does. This suggests that visually independent semantic coding is organized heterogenously along the occipital cortex.

**Highlights:**

- Body-selective lateral occipitotemporal cortex (EBA) contains the strongest visually-independent semantic representations in human visual cortex.
- After removing visual feature variance, semantic encoding is significantly greater in lateral stream regions than in canonical ventral stream areas (FFA, PPA, RSC).The lateral-over-ventral dissociation is architecture-invariant, replicating across six combinations of language models (BERT, GPT-2, CLIP-text) and visual feature sets, with GPT-2 > BERT > CLIP-text ordering validating the pipeline.

## 1. Introduction

The prevailing account of human visual processing describes a representational hierarchy along the ventral stream in which early areas encode local spatial structure while successive higher regions encode increasingly abstract categorical information (Güçlü and van Gerven, 2015). Landmark work mapping semantic selectivity across cortex confirmed that high-level visual areas and adjacent regions encode rich concept-level content aligned with the linguistic semantic system (Contier et al., 2024). Popham et al. subsequently demonstrated that visual and linguistic semantic representations are topographically contiguous at the border of visual cortex: across categories, the pattern of semantic selectivity observed in visually-driven areas sits in immediate correspondence with the pattern in the adjacent language-driven network on the anterior side of the boundary, suggesting the two networks are smoothly joined into a single contiguous map (Popham et al., 2021). What remains unresolved is precisely where semantic content is most concentrated *within* the visual cortex once the contribution of low-level and mid-level visual features is removed—and whether that location corresponds to the region predicted by the canonical ventral hierarchy.

Recent work using large language model (LLM) embeddings of image captions as predictors of fMRI responses has powerfully demonstrated that linguistic-semantic features explain substantial variance in high-level visual cortex. Doerig et al. showed that text-only MPNet embeddings of COCO scene captions successfully predict responses across higher visual areas—including the extrastriate body area (EBA), fusiform face area (FFA), parahippocampal place area (PPA), and retrosplenial complex (RSC)—and that this mapping is sufficiently robust to reconstruct accurate scene captions from brain activity (Doerig et al., 2025). Wang et al. compared language-supervised and vision-only models on the Natural Scenes Dataset (NSD) (Allen et al., 2022), finding that language supervision accounts for greater unique variance specifically in EBA and parts of FFA, suggesting—but not directly testing— that semantic content is disproportionate in lateral body-selective regions (Wang et al., 2023). However, both studies share a critical limitation: neither conditions on low-level visual features. It is therefore impossible to determine from either study whether the observed language-model advantage reflects genuinely unique semantic content, or merely better visual feature approximation by language-supervised models. Convergent evidence from a concurrent preprint using representational similarity analysis on 7T NSD data similarly identifies a lateral occipitotemporal route selective for animate and person-relevant content (Marcos-Manchón and Fuentemilla, 2025), yet that study likewise does not condition on visual features, leaving the attribution of lateral-stream advantage to visually-independent semantic content unresolved. The question of which cortical region retains the most semantic information *after* visual variance is removed cannot be answered by model-comparison or RSA approaches alone.

The methodological framework required to address this question is variance partitioning. Nunez-Elizalde et al. introduced banded ridge regression for jointly fitting multiple correlated feature spaces with separate regularisation per space, showing that naïve joint ridge regression—which uses a single shared regularisation parameter— spuriously inflates semantic predictions in early visual areas due to correlation with low-level visual features, and that banded ridge regression corrects this by assigning independent regularisation per feature space (Nunez-Elizalde et al., 2019). Building on this, Dupré la Tour et al. formalised variance decomposition over large numbers of feature spaces using the product measure, enabling explicit unique-variance attribution across cortex at scale (Dupré la Tour et al., 2022). Lin et al. subsequently applied structured variance partitioning to the NSD using stacked AlexNet encoding models, finding that EBA voxels require only late CNN layers (FC-6 and FC-7) to reach near-maximum prediction accuracy (Lin et al., 2024). This result establishes that EBA tuning is concentrated at the highest levels of a visual feature hierarchy, consistent with abstract categorical representation; however, because all three variance partitioning studies restricted their feature spaces to CNN layers and visual or acoustic features, with no semantic embeddings included, they leave entirely open whether this high-level selectivity is visual-categorical or genuinely linguistic-semantic in nature.

The extrastriate body area occupies an ambiguous position within this debate. Originally identified by Downing et al. as a body-selective region in lateral occipitotemporal cortex (Downing et al., 2001), EBA has traditionally been characterised as a perceptual area encoding the shape and posture of visible bodies without direct involvement in higher-order semantic computation. Peelen and Downing reviewed the functional architecture of body-selective regions, framing EBA as occupying a relatively early role in social vision—a low-to-mid-level perceptual stage rather than a semantic one (Peelen and Downing, 2007). This position was defended by Downing and Peelen (Downing and Peelen, 2011), who reviewed evidence across many proposed higher-level functions for EBA and the fusiform body area (FBA), concluding that these regions encode the shape and posture of perceived bodies but are not implicated in higher-order processes such as identity, emotion, or goal-directed action. One of the most direct empirical tests of this view using a modern encoding model approach is provided by Marrazzo et al., who used ultra-high field 7T fMRI and banded ridge regression with isolated body stimulus images, finding that EBA activity is best explained by a combination of low-level visual and postural features, with no semantic feature space included in their model (Marrazzo et al., 2023).

Converging evidence from multiple paradigms challenges this strictly perceptual account, however. Using representational similarity analysis across person-selective regions of lateral and ventral occipitotemporal cortex, Bracci et al. showed that the *internal organisation* of body part representations is better captured by functional-semantic similarity than by visual shape, physical proximity, or cortical homunculus proximity, suggesting that high-level visual cortex organises body part representations according to their semantic and functional roles (Bracci et al., 2015). An fMRI adaptation study by Kubiak and Króliczak used a repetition suppression paradigm with gesture videos to show that left rostral EBA selectively tracks the meaning of communicative intransitive gestures—exhibiting suppression specific to changes in gesture meaning rather than visual form—indicating higher-order semantic functions of EBA and its involvement in the broader semantic network (Kubiak and Króliczak, 2016). Most recently, Gandolfo et al. provided causal evidence, using a large-scale fMRI re-analysis (*n* = 92) and fMRI-guided transcranial magnetic stimulation, that the left EBA responds preferentially to face-to-face social dyads and that online TMS over left EBA specifically abolishes the behavioural marker of visual sensitivity to social interactions—demonstrating that this region causally supports social action perception rather than merely visual body detection (Gandolfo et al., 2024). These findings collectively suggest that EBA participates in higher-order semantic and social computations, yet no prior study has directly quantified the unique semantic information EBA carries beyond visual feature predictions using a naturalistic stimulus set, and no systematic cross-ROI comparison on a common unique-semantic-information metric has been conducted.

The present study closes this gap by asking a question that no prior work has directly addressed: after conditioning on a comprehensive low-to-mid-level visual feature hierarchy, which visual ROI retains the greatest visually-independent variance explained by language model embeddings of COCO image captions? We apply cross-validated ridge regression to the NSD (Allen et al., 2022), residualising language model embeddings against a visual feature hierarchy before using them to predict voxelwise brain responses—thereby isolating semantic content that cannot be attributed to the visual features we modelled. We find that visually-independent semantic information peaks in EBA, not in FFA, PPA, or RSC, and that This lateral-over-ventral dissociation is consistent across all eight subjects and both hemispheres. Early visual cortex shows robustly negative residual predictions, providing an internal negative control that confirms the residualisation selectively removes visually-driven signal rather than amplifying it. The dissociation is replicated across six combinations of language models (BERT, GPT-2, CLIP-text) and visual feature architectures, ruling out model-specific confounds.

This result directly reframes the finding of Lin et al. (Lin et al., 2024) that EBA requires only late CNN layers: where that analysis concluded from CNN-only partitioning that EBA tuning is concentrated at high visual abstraction levels, the addition of a linguistic feature space reveals that this high-level selectivity is semantic rather than purely visual-categorical in nature. Because both analyses use variance partitioning on the NSD, the comparison is direct—the sole difference is the absence of a linguistic feature space in Lin et al. (Lin et al., 2024), which is precisely what concealed EBA’s semantic profile. The finding equally challenges the characterisation of EBA as a cognitively unelaborated perceptual detector (Downing and Peelen, 2011), and stands in contrast to Marrazzo et al. (Marrazzo et al., 2023), whose null semantic result we suggest reflects the restricted semantic range of an isolated body-stimulus design rather than a genuine absence of semantic content in EBA under naturalistic viewing conditions. Our approach extends the topographic observation of Popham et al. (Popham et al., 2021) from a claim about spatial adjacency to a claim about representational content: EBA does not merely neighbour the linguistic semantic system; its own visually-evoked responses carry semantic information that is orthogonal to visual features. This is consistent with its causal role in social interaction perception (Gandolfo et al., 2024) and with prior evidence for semantic organisation of body part representations in lateral occipitotemporal cortex (Bracci et al., 2015; Kubiak and Króliczak, 2016). Unlike LLM-based prediction studies that establish broad alignment between language models and higher visual cortex without conditioning on visual features (Doerig et al., 2025; Wang et al., 2023), our residualisation approach reveals a clear ordering among visual ROIs that is invisible without explicit visual conditioning. We conclude that the human visual system encodes not only what objects look like, but carries semantic structure extending beyond visual features, and that this capacity is concentrated in body-selective lateral occipitotemporal cortex—consistent with its established roles in action and social perception (Gandolfo et al., 2024; Bracci et al., 2015; Kubiak and Króliczak, 2016).

## 2. Methods

### 2.1. Dataset

We used data from the Natural Scenes Dataset (NSD; (Allen et al., 2022)), a large-scale 7T fMRI dataset in which participants viewed tens of thousands of colour photographs drawn from the MS-COCO image corpus (Lin et al., 2014). We analysed the data for eight subjects (subj01–subj08), using the preprocessed fMRI responses in challenge space (z-scored betas, one response per trial per voxel) for both left (LH) and right (RH) hemispheres. Region-of-interest (ROI) masks were taken from the NSD challenge-space parcellations across six functional localiser types: early retinotopic areas (`prf-visualrois`: V1v, V1d, V2v, V2d, V3v, V3d, hV4), body-selective areas (`floc-bodies`: EBA, FBA-1, FBA-2, mTL-bodies), face-selective areas (`floc-faces`: OFA, FFA-1, FFA-2, mTL-faces, aTL-faces), placeselective areas (`floc-places`: OPA, PPA, RSC), word-selective areas (`floc-words`: OWFA, VWFA-1, VWFA-2, mfs-words, mTL-words), and stream-level summary ROIs (`streams`: early, midventral, midlateral, midparietal, ventral, lateral, parietal). For each trial, the corresponding MS-COCO image ID was retrieved via the NSD stimulus information file (`nsd_stim_info_merged.pkl`), and all available COCO category labels and captions for that image were concatenated into a single text string used as the semantic descriptor for that trial.

### 2.2. Visual Feature Extraction

Two complementary visual feature sets were extracted for each training image.

#### L1: Gabor filterbank

A bank of 8 × 4 = 32 Gabor filters was constructed by crossing 8 orientations (*θ* = *kπ/*8, *k* = 0, …, 7) with 4 spatial scales (*σ* = 2^*s*^, *λ* = 2^*s*+1^, *s* = 0, …, 3; kernel size 21 × 21 pixels). Each filter response was summarised by its mean and standard deviation computed within a multi-scale spatial grid (1 × 1, 2 × 2, and 4 × 4 non-overlapping cells), yielding a compact energy representation of local orientation content across the image. This was supplemented by HSV colour histograms (16 bins per channel, in a 2 × 2 spatial grid) and Canny edge density (3 threshold pairs, 2 × 2 grid). All Gabor features were computed on single-channel (greyscale) versions of the image; colour and edge features were appended to form the complete L1 vector.

#### L2: VGG19 deep visual features

Images were resized to 224 × 224 pixels and passed through a pretrained VGG19 network (Simonyan and Zisserman, 2015) (ImageNet weights). The output of the global average pooling layer following the final convolutional block (`avgpool`, 512 dimensions) was used as the full-dimensionality L2 feature vector. No dimensionality reduction was applied.

The combined visual feature matrix 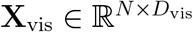 was formed by horizontal concatenation of the L1 Gabor vector and the L2 VGG19 vector. All feature extraction was performed with no PCA or other compression at any stage.

### 2.3. Language Model Feature Extraction

Semantic features were extracted using BERT (`bert-base-uncased`; (Devlin et al., 2019)). The concatenated COCO category and caption string for each trial was tokenised (maximum length 512 tokens, padding and truncation applied) and passed through the pretrained BERT encoder. The `[CLS]` token representation from the final hidden layer was used as the trial-level semantic embedding, yielding a 768-dimensional vector **b**_*i*_ ∈ ℝ^768^ for each trial *i*.

### 2.4. Visual Residualisation of Language Embeddings

To isolate semantic information in BERT embeddings that is not linearly predictable from visual features, we applied a cross-validated residualisation procedure.

#### Alpha selection

A RidgeCV regression (*α* ∈ {10^1^, 10^1.45^, …, 10^6^}, 12 log-spaced values; 5-fold internal cross-validation; scoring by mean *R*^2^ across BERT dimensions) was fit once on the full standardised **X**_vis_ for each subject to select a single regularisation parameter *α*^∗^. The input was standardised (zero mean, unit variance per feature) before regression. The selected *α*^∗^ was held fixed for all subsequent steps within that subject. The selected *α*^∗^ was identical across all eight subjects (*α*^∗^ = 123,284.67), indicating that the linear relationship between the visual feature space and BERT embeddings is highly consistent across individuals.

#### Cross-validated residuals

A 5-fold KFold split (shuffled, fixed random seed) was applied to the *N* trials. For each fold, a `StandardScaler` was fit exclusively on the training fold of **X**_vis_ and applied to both training and test folds (preventing data leakage). A Ridge regression with the selected *α*^∗^ was then fit to predict all 768 BERT dimensions simultaneously from the scaled training-fold visual features. The out-of-fold BERT residual was defined as:

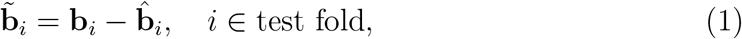

where 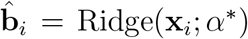 is the visual prediction of BERT for trial *i*. Out-of-fold prediction was used throughout to prevent in-sample overfitting from artificially inflating the amount of visual variance removed. The complete residualised embedding matrix 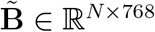 was assembled by concatenating held-out fold residuals. By construction, 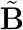 is orthogonal in the linear sense to the visual feature subspace; this orthogonality is the basis for the mechanistically interpretable negative 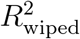 in early visual cortex, where residualised embeddings anti-predict voxel responses that are dominated by visual structure. We note that this residualisation assumes a linear relationship between visual and semantic feature spaces; to the extent that nonlinear dependencies exist between deep network features and language model embeddings, the residual may retain a small visually-related component, a limitation shared by banded ridge regression approaches.

#### Rationale for sequential residualisation

The sequential residualisation operates in feature space rather than response space, which distinguishes it from banded ridge regression (BRR) and makes it a more direct operationalization of visually-independent semantic content. Our approach instead removes visually-predictable variance from the language model embeddings themselves before they enter any brain encoding model, ensuring the residual is geometrically orthogonal to the visual feature subspace by construction. These two approaches answer subtly different questions: BRR quantifies the unique brain variance attributable to semantics, while our residualisation isolates the component of the semantic representation that is irrecoverable from visual features and asks whether that component predicts neural responses — a more direct operationalisation of visually-independent semantic content. A further practical advantage is that the residual’s orthogonality to the visual subspace produces the mechanistically interpretable negative predictions in early visual cortex that serve as an internal validity check, a result unavailable under BRR where unique variance estimates are bounded below at zero. Because regularisation in the encoding step shrinks estimates toward zero rather than inflating them, any bias from the fixed *α* = 10^4^ makes positive findings in higher visual areas conservative underestimates rather than inflations; the consistency of the regional dissociation across six architecture combinations provides further assurance that results are not an artefact of this choice. The two approaches answer genuinely different scientific questions. BRR asks: after partialling, how much unique brain variance does semantics explain? The residualisation asks: does the part of the language model that is geometrically orthogonal to vision predict brain responses? These are not equivalent. The first is a property of the brain response; the second is a property of the representation itself.

### 2.5. Voxelwise Encoding Model

For each subject and hemisphere, a voxelwise encoding model was fit to predict fMRI responses from the residualised BERT embeddings 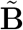 using 5-fold cross-validated Ridge regression (*α* = 10^4^, fixed). Voxels were processed in parallel blocks of 5,000. For each fold, 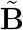 was standardised on the training set (zero mean, unit variance per dimension) before fitting; the same scaler was applied to the test set. The cross-validated *R*^2^ for each voxel *v* was computed as:

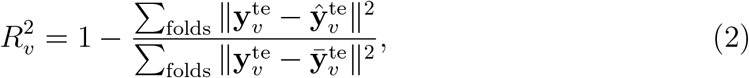

where sums of squares accumulate across all held-out folds before the ratio is taken (avoiding instability in small folds). This quantity, 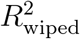, represents variance in the fMRI signal explained by semantic content that is not shared with the visual features. For comparison, an identical procedure was applied using the raw (non-residualised) BERT embeddings **B** to obtain 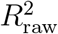. See Figure 1

**Figure 1.**
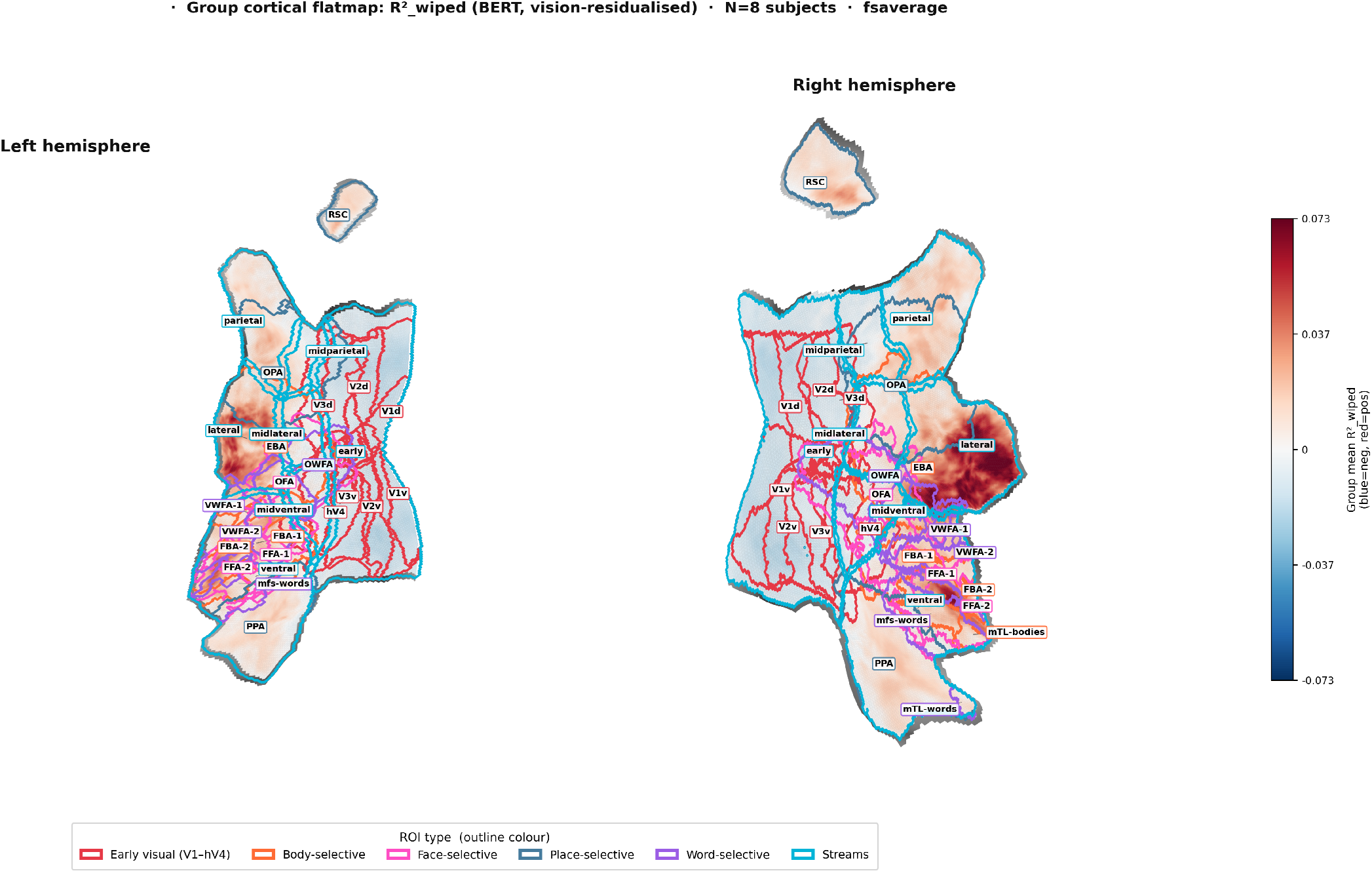
Group-level cortical flatmap of vision-residualised BERT representations across the ventral and dorsal visual hierarchy. Colour indicates group-mean R^2^wiped — the variance in fMRI responses (N = 8 subjects) explained by BERT text embeddings after removing shared variance with a visual feature model (VGG-19). Red indicates positive unique BERT variance; blue indicates regions where BERT explains less variance than vision alone (i.e., negative residual, as expected in early retinotopic cortex). Coloured outlines demarcate individual ROIs from six functional atlas systems: early visual cortex (V1v–hV4; red), body-selective regions (EBA, FBA-1/2, mTL-bodies; orange), face-selective regions (OFA, FFA-1/2, mTL/aTL-faces; pink), place-selective regions (OPA, PPA, RSC; blue), word-selective regions (OWFA, VWFA-1/2, mfs/mTL-words; purple), and visual streams (early, midventral/lateral/parietal, ventral, lateral, parietal; cyan). Data are projected onto the fsaverage flat surface. Left and right hemisphere flatmaps are shown separately; RSC appears as a spatially isolated island consistent with its medial-wall position on the unfolded cortex. Colour scale is symmetric around zero (vmax = 99th percentile of positive vertices across both hemispheres).

### 2.6. ROI Aggregation

For each ROI, 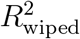 was averaged across all voxels within the ROI mask to obtain a single scalar per subject per hemisphere. Only ROIs with at least 5 contributing voxels were retained. The semantic independence ratio was defined as 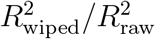 expressing the proportion of total explainable BERT variance that survives visual partialling. As an additional voxel-level diagnostic, the fraction of voxels within each ROI with 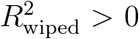 (`frac_pos_wiped`) was recorded per subject and used to characterise the sign of residual predictions across the cortical hierarchy.

### 2.7. Statistical Analysis

Group-level inference treated subjects as the random effect. For each ROI and hemisphere, a one-sample *t*-test was performed against zero on the vector of persubject 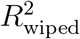 means. Effect sizes were quantified using Hedges’ *g*:

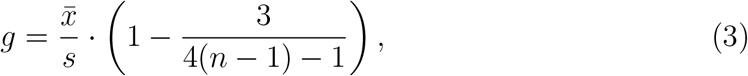

where 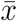 and *s* are the sample mean and standard deviation across subjects and *n* is the number of subjects. Multiple comparisons across all ROI × hemisphere tests were controlled using the Benjamini–Hochberg false discovery rate (BH-FDR) procedure at *q <* 0.05. To confirm significant results without distributional assumptions, we additionally applied exact sign-flip permutation tests: all 2^*N*^ sign combinations (exhaustive with *N* = 8 subjects, giving 256 permutations) were evaluated, and the permutation *p*-value was defined as the proportion of permuted group means ≥ the observed mean. Sign-flip *p*-values were also BH-FDR corrected across all tests. All analyses were implemented in Python using `scikit-learn` (Pedregosa et al., 2011), `scipy` (Virtanen et al., 2020), and `statsmodels` (Seabold and Perktold, 2010).

This study used a pre-existing public dataset (NSD). ROI definitions were taken from the NSD challenge-space parcellations without modification. The primary hypothesis — that lateral occipitotemporal cortex would show greater visually-independent semantic encoding than ventral stream regions — was pre-specified based on prior literature. The VWFA-2 finding was not predicted in advance and should be treated as exploratory.

### 2.8. Architecture Robustness Analysis

To assess whether the observed encoding effects depended on the specific choice of language model or visual feature architecture, the full residualisation and voxelwise prediction pipeline was repeated across a 3 × 2 factorial design, yielding six independent wipe conditions per subject per ROI.

#### Language models

Three pretrained language models were evaluated: (1) BERT-base (`bert-base-uncased`; (Devlin et al., 2019)), using the `[CLS]` token representation from the final hidden layer (768 dimensions); (2) GPT-2 (`gpt2`; (Radford et al., 2019)), using the last-token hidden state from the final transformer block (768 dimensions); and (3) CLIP-text (`openai/clip-vit-base-patch32`; (Radford et al., 2021)), using the projected text embedding (512 dimensions). For all three models, each trial’s text descriptor was constructed by concatenating the MS-COCO category labels and captions associated with the viewed image, as described above.

#### Visual feature variants

Two visual feature matrices were constructed. Variant **X**_*a*_ was a broad ensemble formed by concatenating five feature sets: the L1 Gabor filterbank, VGG19 global average pooling output (512 dimensions), ResNet50 global average pooling output (2,048 dimensions; (He et al., 2016)), CLIP-visual image embeddings (512 dimensions), and EfficientNet-B0 global average pooling output (1,280 dimensions; (Tan and Le, 2019)), for a combined dimensionality of 4,352. Variant **X**_*b*_ was a VGG19 hierarchical multi-layer feature set obtained by applying global average pooling to the relu3_4, relu4_4, and relu5_4 activation maps (256, 512, and 512 dimensions respectively), concatenated to 1,280 dimensions. The motivation for **X**_*a*_ was maximal coverage of the visual feature space; the motivation for **X**_*b*_ was a controlled, architecture-homogeneous baseline using only hierarchical VGG19 activations.

#### Wipe conditions

Crossing the three language models with the two visual feature variants produced six wipe conditions: `bert_Xa`, `bert_Xb`, `gpt2_Xa`, `gpt2_Xb`, `clip_Xa`, and `clip_Xb`. For each condition, the residualisation procedure was applied independently using a single 80/20 train/test split in place of 5-fold cross-validation, chosen for computational tractability across six model combinations per subject. RidgeCV alpha selection was performed on the training 80% of trials; the language model embedding residual was computed on the held-out test 20% only, and voxelwise Ridge encoding models (*α* = 10^4^, fixed) were fit exclusively on these test-set trials. This procedure yields conservative 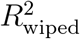 estimates relative to the primary analysis owing to the reduced trial count; absolute values in the robustness heatmap are therefore not directly comparable to primary analysis estimates, and comparisons are made within the heatmap across conditions rather than against the primary analysis figures. Each condition therefore produced its own set of per-subject, per-voxel 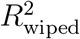 maps, and its own ROI aggregations entering the group-level statistical comparisons described above. The **X**_*a*_ conditions remove more visual variance from the language embedding by virtue of the broader feature ensemble, leaving a smaller but cleaner residual; the **X**_*b*_ conditions provide a more conservative benchmark. Convergence of the regional dissociation pattern across all six conditions was taken as evidence of architecture-independent robustness.

## 3. Results

### 3.1. Visual features capture a substantial but incomplete portion of language model variance

Before testing whether non-visual semantic information predicts brain responses, we quantified how much variance in BERT representations could be accounted for by visual features alone. Across all eight subjects, the visual feature set (Gabor filterbank concatenated with full VGG19-avgpool features) explained a consistent 42.9–43.2% of variance in BERT embeddings (mean 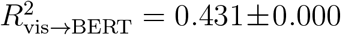 across subjects; RidgeCV-selected regularisation *α* = 123, 284.67, identical across all eight subjects). The remaining ~ 57% of BERT variance was orthogonal to the visual feature space and constituted the residual signal tested in all subsequent analyses. This partial but substantial overlap confirms that visual and semantic features share a meaningful subspace, motivating the cross-validated residualisation as a necessary preprocessing step before any claims about non-visual semantic coding can be made.

### 3.2. Non-visual semantic signals are detectable across higher visual cortex

We tested whether residualised BERT embeddings—from which all predictable visual variance had been removed via held-out 80/20 split regression—explained significant variance in voxelwise fMRI responses. Across 55 region-of-interest tests in 8 subjects, 40 ROIs (73%) survived BH-FDR correction at *q <* 0.05 by random-effects one-sample *t*-test, with subjects treated as the unit of inference. Of these, 24 were additionally confirmed by exact sign-flip permutation test (2^8^ = 256 exhaustive permutations, BH-FDR corrected), which makes no distributional assumptions. The 16 ROIs significant by *t*-test but not sign-flip were predominantly lower-effect-size regions for which *n <* 8 prevents the sign-flip test from reaching sufficient resolution (minimum achievable *p* = 1*/*2^*n*^).

The strongest effects were concentrated in lateral occipitotemporal cortex. Lateral stream ROI reached 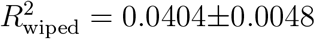 SEM (RH) and 0.0290*±*0.0035 SEM (LH), with Hedges *g* = 2.62 and 2.64 respectively (both *q*_FDR_ *<* 0.0003, signflip *p* = 0.0039, the minimum achievable with *N* = 8). EBA reached 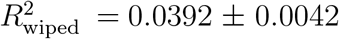 (RH) and 0.0282 ± 0.0037 (LH), with Hedges *g* = 2.90 and 2.41 (both *q*_FDR_ *<* 0.0004, sign-flip *p* = 0.0039). These effect sizes are large by any standard in encoding model work. The body-selective FBA-2 showed consistent effects bilaterally (RH: 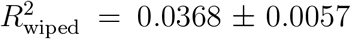, *g* = 2.02, *q*_FDR_ = 0.001; LH: 0.0332 ± 0.0051, *g* = 2.14, *q*_FDR_ = 0.001), confirming the result is not isolated to a single body-selective region. Face-selective cortex showed smaller but reliable effects: FFA-1 reached 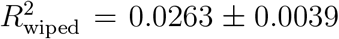 (RH, *g* = 2.10, *q*_FDR_ = 0.001) and 0.0189 *±* 0.0042 (LH, *g* = 1.40, *q*_FDR_ = 0.005). Left VWFA-2 showed significant non-visual semantic prediction (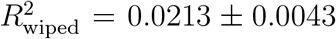, *g* = 1.55, *q*_FDR_ = 0.004, sign-flip *q* = 0.019, *N* = 8). The right hemisphere showed a numerically positive effect (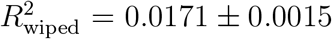, *g* = 4.11) that survived *t*-test FDR correction but not sign-flip (*q*_signflip_ = 0.061), and should be treated with caution: only *N* = 5 subjects had adequate RH VWFA-2 coverage, making this finding underpowered for the present sample. The left-hemisphere result is the interpretable one.

(See: regional dissociation bar plot with all ROIs.Figure 2 panel A and table 1)

**Table 1:** Group-level visually-independent semantic encoding 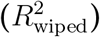 across visual ROIs. Values are mean *±* SEM across subjects. 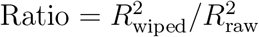. *g* = Hedges’ *g* (effect size). *q*_FDℝ_: BH-corrected *p*-value from one-sample *t*-test. *q*_SF_: BH-corrected sign-flip permutation *p*-value (2^*N*^ exhaustive permutations). Rows are sorted by descending 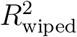 within each ROI category.

| ROI | H | N | $R^2_{\text{wiped}}$ (mean ± SEM) | $R^2_{\text{raw}}$ | Ratio | $g$ | $q_{\text{FDR}}$ | $q_{\text{SF}}$ |
| --- | --- | --- | --- | --- | --- | --- | --- | --- |
| *Stream-level ROIs* |  |  |  |  |  |  |  |  |
| Lateral stream | RH | 8 | 0.0404 ± 0.0048 | 0.219 | 0.181 | 2.62 | <0.001 | 0.020* |
| Lateral stream | LH | 8 | 0.0290 ± 0.0035 | 0.175 | 0.167 | 2.64 | <0.001 | 0.020* |
| Ventral stream | RH | 8 | 0.0138 ± 0.0015 | 0.161 | 0.085 | 2.87 | <0.001 | 0.020* |
| Ventral stream | LH | 8 | 0.0125 ± 0.0026 | 0.149 | 0.081 | 1.53 | 0.004 | 0.020* |
| Parietal stream | RH | 8 | 0.0120 ± 0.0027 | 0.103 | 0.113 | 1.38 | 0.006 | 0.020* |
| Parietal stream | LH | 8 | 0.0098 ± 0.0020 | 0.105 | 0.087 | 1.53 | 0.004 | 0.020* |
| Mid-lateral | RH | 8 | 0.0046 ± 0.0028 | 0.103 | 0.022 | 0.51 | 0.178 | 0.136 |
| Mid-lateral | LH | 8 | 0.0035 ± 0.0025 | 0.094 | 0.029 | 0.45 | 0.223 | 0.208 |
| Mid-parietal | LH | 8 | 0.0007 ± 0.0027 | 0.098 | −0.013 | 0.08 | 0.850 | 0.767 |
| Mid-parietal | RH | 8 | −0.0001 ± 0.0019 | 0.085 | −0.011 | −0.02 | 0.963 | 0.772 |
| Mid-ventral | RH | 8 | −0.0008 ± 0.0028 | 0.083 | −0.044 | −0.09 | 0.850 | 0.876 |
| Mid-ventral | LH | 8 | −0.0020 ± 0.0025 | 0.088 | −0.044 | −0.25 | 0.505 | 1.000 |
| Early stream | RH | 8 | −0.0104 ± 0.0007 | 0.047 | −0.242 | −4.66 | <0.001 | 1.000 |
| Early stream | LH | 8 | −0.0107 ± 0.0007 | 0.048 | −0.241 | −5.02 | <0.001 | 1.000 |
| *Body-selective* |  |  |  |  |  |  |  |  |
| EBA | RH | 8 | 0.0392 ± 0.0042 | 0.222 | 0.174 | 2.90 | <0.001 | 0.020* |
| FBA-2 | RH | 8 | 0.0368 ± 0.0057 | 0.238 | 0.150 | 2.02 | <0.001 | 0.020* |
| FBA-2 | LH | 7 | 0.0332 ± 0.0051 | 0.180 | 0.192 | 2.14 | 0.001 | 0.020* |
| EBA | LH | 8 | 0.0282 ± 0.0037 | 0.176 | 0.158 | 2.41 | <0.001 | 0.020* |
| FBA-1 | LH | 5 | 0.0263 ± 0.0045 | 0.151 | 0.183 | 2.09 | 0.006 | 0.061<sup>†</sup> |
| FBA-1 | RH | 6 | 0.0127 ± 0.0039 | 0.140 | 0.085 | 1.11 | 0.033 | 0.061<sup>†</sup> |
| mTL-bodies | RH | 2 | 0.0060 ± 0.0008 | 0.050 | 0.128 | 0.00 | 0.109 | 0.417 |
| *Face-selective* |  |  |  |  |  |  |  |  |
| FFA-1 | RH | 8 | 0.0263 ± 0.0039 | 0.201 | 0.126 | 2.10 | <0.001 | 0.020* |
| FFA-2 | LH | 7 | 0.0233 ± 0.0078 | 0.154 | 0.109 | 0.98 | 0.034 | 0.036* |

Table 1 – *continued from previous page*
| ROI | H | N | $R^2_{\text{wiped}}$ (mean ± SEM) | $R^2_{\text{raw}}$ | Ratio | $g$ | $q_{\text{FDR}}$ | $q_{\text{SF}}$ |
| --- | --- | --- | --- | --- | --- | --- | --- | --- |
| FFA-2 | RH | 8 | $0.0213 \pm 0.0051$ | 0.168 | 0.087 | 1.31 | 0.006 | 0.020* |
| FFA-1 | LH | 8 | $0.0189 \pm 0.0042$ | 0.168 | 0.104 | 1.40 | 0.005 | 0.020* |
| OFA | LH | 8 | $0.0067 \pm 0.0038$ | 0.120 | 0.030 | 0.55 | 0.147 | 0.111 |
| OFA | RH | 8 | $0.0051 \pm 0.0038$ | 0.099 | 0.005 | 0.43 | 0.242 | 0.208 |
| *Place-selective* |  |  |  |  |  |  |  |  |
| RSC | LH | 8 | $0.0105 \pm 0.0037$ | 0.172 | 0.051 | 0.89 | 0.036 | 0.020* |
| RSC | RH | 8 | $0.0103 \pm 0.0022$ | 0.168 | 0.059 | 1.48 | 0.004 | 0.020* |
| PPA | LH | 8 | $0.0103 \pm 0.0024$ | 0.190 | 0.051 | 1.34 | 0.006 | 0.020* |
| OPA | RH | 8 | $0.0102 \pm 0.0021$ | 0.133 | 0.074 | 1.52 | 0.004 | 0.020* |
| PPA | RH | 8 | $0.0100 \pm 0.0020$ | 0.192 | 0.049 | 1.56 | 0.004 | 0.020* |
| OPA | LH | 8 | $0.0094 \pm 0.0022$ | 0.140 | 0.062 | 1.36 | 0.006 | 0.020* |
| *Word-selective* |  |  |  |  |  |  |  |  |
| VWFA-2 | LH | 8 | $0.0213 \pm 0.0043$ | 0.114 | 0.172 | 1.55 | 0.004 | 0.020* |
| VWFA-1 | RH | 8 | $0.0181 \pm 0.0021$ | 0.156 | 0.118 | 2.66 | <0.001 | 0.020* |
| VWFA-2 | RH | 5 | $0.0171 \pm 0.0015$ | 0.141 | 0.124 | 4.11 | <0.001 | 0.061<sup>†</sup> |
| VWFA-1 | LH | 8 | $0.0126 \pm 0.0029$ | 0.122 | 0.100 | 1.35 | 0.006 | 0.020* |
| mfs-words | LH | 8 | $0.0119 \pm 0.0051$ | 0.127 | 0.072 | 0.74 | 0.066 | 0.019 |
| mfs-words | RH | 6 | $0.0104 \pm 0.0039$ | 0.122 | 0.074 | 0.93 | 0.058 | 0.061 |
| OWFA | LH | 8 | $0.0004 \pm 0.0026$ | 0.070 | -0.036 | 0.04 | 0.929 | 0.733 |
| OWFA | RH | 8 | $-0.0002 \pm 0.0029$ | 0.061 | -0.074 | -0.02 | 0.963 | 0.772 |
| *Early visual (retinotopic)* |  |  |  |  |  |  |  |  |
| hV4 | LH | 8 | $-0.0036 \pm 0.0020$ | 0.079 | -0.061 | -0.57 | 0.139 | 1.000 |
| hV4 | RH | 8 | $-0.0041 \pm 0.0017$ | 0.070 | -0.067 | -0.76 | 0.061 | 1.000 |
| V3v | RH | 8 | $-0.0078 \pm 0.0013$ | 0.057 | -0.148 | -1.83 | 0.001 | 1.000 |
| V3d | LH | 8 | $-0.0079 \pm 0.0013$ | 0.050 | -0.181 | -1.95 | 0.001 | 1.000 |
| V3d | RH | 8 | $-0.0080 \pm 0.0009$ | 0.052 | -0.168 | -2.68 | <0.001 | 1.000 |
| V3v | LH | 8 | $-0.0082 \pm 0.0013$ | 0.061 | -0.164 | -1.94 | 0.001 | 1.000 |
| V2d | LH | 8 | $-0.0105 \pm 0.0009$ | 0.048 | -0.246 | -3.56 | <0.001 | 1.000 |
| V2d | RH | 8 | $-0.0107 \pm 0.0007$ | 0.051 | -0.227 | -5.01 | <0.001 | 1.000 |
| V2v | RH | 8 | $-0.0109 \pm 0.0009$ | 0.060 | -0.220 | -3.99 | <0.001 | 1.000 |
| V2v | LH | 8 | $-0.0121 \pm 0.0007$ | 0.067 | -0.206 | -5.71 | <0.001 | 1.000 |
| V1d | LH | 8 | $-0.0138 \pm 0.0008$ | 0.065 | -0.231 | -5.29 | <0.001 | 1.000 |
| V1v | RH | 8 | $-0.0140 \pm 0.0007$ | 0.063 | -0.257 | -6.16 | <0.001 | 1.000 |

Table 1 – *continued from previous page*
| ROI | H | N | $R^2_{\text{wiped}}$ (mean ± SEM) | $R^2_{\text{raw}}$ | Ratio | $g$ | $q_{\text{FDR}}$ | $q_{\text{SF}}$ |
| --- | --- | --- | --- | --- | --- | --- | --- | --- |
| V1d | RH | 8 | $-0.0141 \pm 0.0005$ | 0.064 | -0.240 | -8.51 | <0.001 | 1.000 |
| V1v | LH | 8 | $-0.0146 \pm 0.0005$ | 0.060 | -0.257 | -8.99 | <0.001 | 1.000 |

H = hemisphere; N = number of subjects contributing to that ROI.
\* sign-flip BH-FDR $q < 0.05$ (confirmed without distributional assumptions).
† significant by $t$ -test FDR only; sign-flip test did not reach threshold, attributable to $N < 7$ (minimum achievable $p = 1/2^N$ ).
All $q_{\text{FDR}}$ values apply BH correction across all $55 \text{ ROI} \times \text{hemisphere}$ tests.

**Figure 2.**
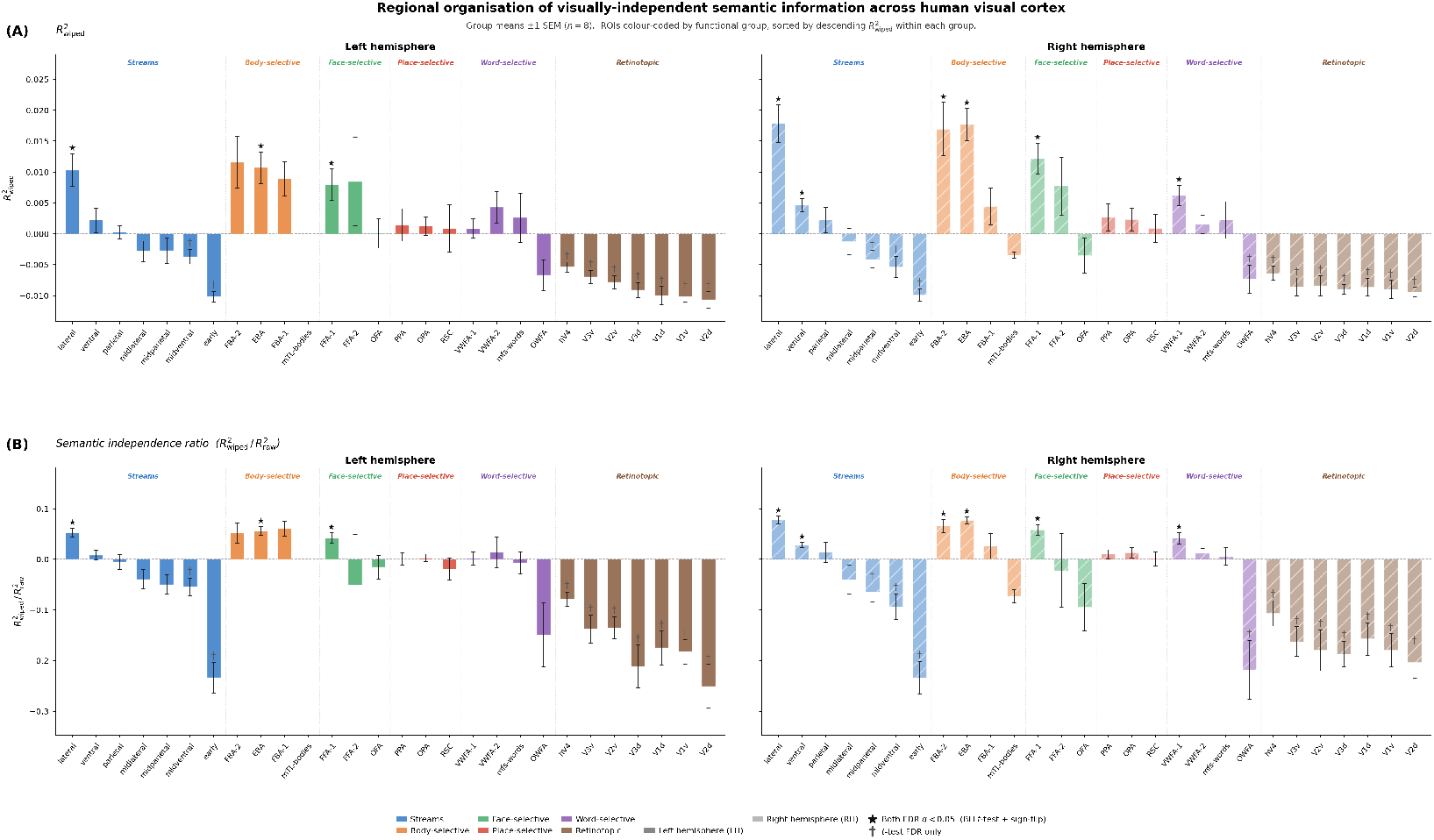
Regional organisation of visually-independent semantic information across human visual cortex. Group means ± 1 SEM (*n* = 8). ROIs colour-coded by functional group (blue = streams; orange = body-selective; green = face-selective; red = place-selective; purple = word-selective; brown = retinotopic), sorted by descending 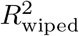 within each group. LH and RH shown in separate columns. **(A)** 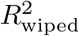 Cross-validated *R*^2^ of voxelwise fMRI responses predicted by visually-residualised BERT embeddings (Gabor + VGG19 visual features removed via 5-fold ridge regression). Positive values indicate semantic variance beyond visual features; negative values indicate anti-prediction. EBA, FBA-2, and lateral stream show the largest positive effects bilaterally; early retinotopic areas (V1–V3, early stream) are uniformly negative, serving as an internal negative control. **(B) Semantic independence ratio** 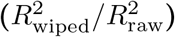. Fraction of total BERT alignment surviving visual partialling. EBA and lateral stream: ≈0.16–0.18; ventral stream and place-selective regions: ≈0.05–0.09 (3–4*×* lower). LH VWFA-2 reaches ≈0.17, discussed in Section 4.6. ⋆ both BH-FDR *t*-test and sign-flip significant (2^8^ permutations, *q <* 0.05); † *t*-test FDR only. All corrections applied jointly across 55 ROI *×* hemisphere tests.

### 3.3. Non-visual semantic coding is systematically stronger in lateral than ventral cortex

A key structural prediction was that lateral occipitotemporal cortex, implicated in social and action-relevant body processing, would show disproportionately stronger non-visual semantic coding than ventral stream regions organised around visual category geometry. This prediction was confirmed bilaterally with consistent magnitude.

In the right hemisphere, lateral stream 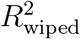 (0.0404) exceeded ventral stream 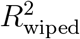 (0.0138) by a factor of 2.93, and EBA (0.0392) exceeded PPA (0.0100) by a factor of 3.91. In the left hemisphere, lateral (0.0290) exceeded ventral (0.0125) by a factor of 2.33, and EBA (0.0282) exceeded PPA (0.0103) by a factor of 2.74. This dissociation was not simply a difference in absolute *R*^2^—it was preserved when expressed as a semantic independence ratio 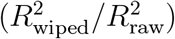, which normalises for differences in overall response magnitude. The ratio for lateral stream was 0.181 (RH) and 0.167 (LH), versus 0.085 (RH) and 0.081 (LH) for ventral stream. The ratio for EBA was 0.174 (RH) and 0.158 (LH), versus 0.049 (RH) and 0.051 (LH) for PPA. This means that approximately 17% of the total explainable variance in EBA is non-visual semantic, compared to approximately 5% in PPA—a 3–4 × difference in the fraction of semantic coding that is visually independent. Ventral stream and place-selective regions did retain significant non-visual semantic signals (ventral bilateral: both *q*_FDR_ *<* 0.004, sign-flip confirmed; PPA bilateral: *q*_FDR_ *<* 0.006, sign-flip confirmed), but at substantially lower magnitudes. The lateral-over-ventral dissociation was fully bilateral, ruling out a hemisphere-specific artefact.Refer Figure 3.

**Figure 3.**
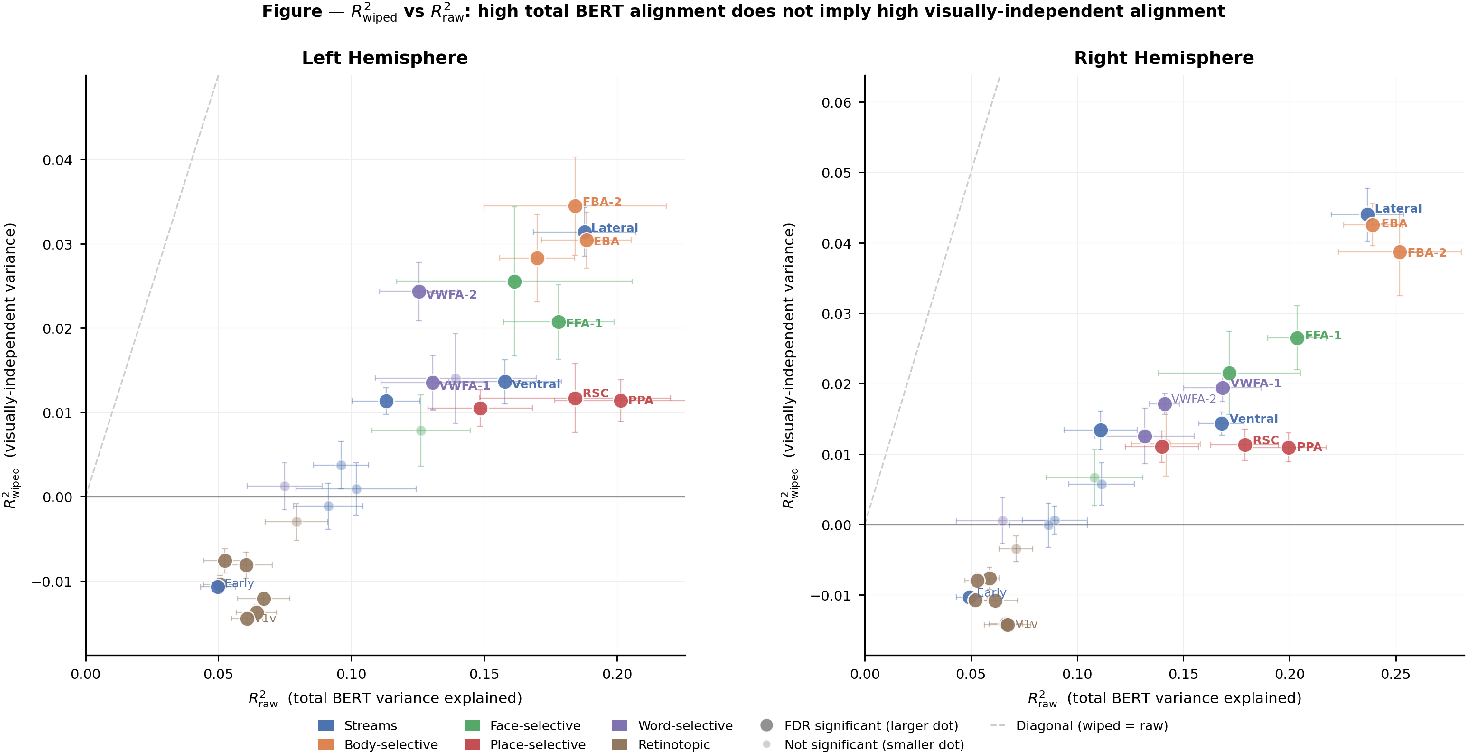
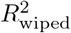 does not scale with 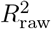 across visual cortex. Each point represents one ROI × hemisphere (LH: left panel; RH: right panel), colour-coded by functional group (± 1 SEM error bars; *n* = 8 subjects). Larger dots passed BH-FDR correction; smaller transparent dots did not. The dashed diagonal marks 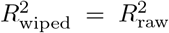 points below zero are anti-predicted by residualised BERT embeddings. Place-selective and ventral regions (PPA, RSC, ventral stream) have high 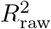 but fall far below the diagonal, indicating their BERT alignment is largely visual in origin. EBA and lateral stream show the highest 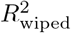 despite moderate 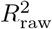. Early retinotopic areas (V1–V3, early stream) fall below zero, serving as an internal negative control. Bold labels indicate significance on both BH-FDR *t*-test and exact sign-flip permutation test (2^8^ permutations, *q <* 0.05).

Left VWFA-2 showed a semantic independence ratio of 0.172, comparable in magnitude to RH EBA (0.174), representing a secondary observation: left word-selective cortex retains nearly 17% of its explainable variance as non-visual semantic content after visual partialling. This finding is specific to the left hemisphere and sign-flip confirmed; the right hemisphere analogue is not robust at the present sample size.

### 3.4. Early visual cortex shows a robust negative signal, validating the residualisation procedure

An important control result was not in the higher cortex but in early visual regions. V1v showed 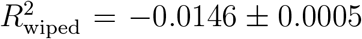 SEM (LH) and −0.0140 *±* 0.0007 (RH), with Hedges *g* = −8.99 and −6.16 respectively—the largest effect sizes in the entire dataset in absolute terms, in the negative direction (both *q*_FDR_ *<* 0.000005). V1d reached *g* = −5.29 (LH) and −8.51 (RH). The early stream summary ROI showed *g* = −5.02 (LH) and −4.66 (RH), both *q*_FDR_ *<* 0.00002. The complete V1–V3 hierarchy was significantly negative bilaterally after FDR correction.

Critically, the sign-flip test for all early visual and V1–V3 ROIs returned *p*_signflip_ = 1.0—meaning every single one of the 256 sign permutations produced a mean equal to or larger than the observed mean. Not one permutation produced a positive group mean in any early visual ROI. At the subject level, `frac_pos_wiped` in V1v was 0.000 for all 8 subjects in LH, and for 6 of 8 subjects in RH (the remaining two were *<* 0.004). In contrast, `frac_pos_wiped` in EBA ranged from 0.718 to 0.959 across subjects and hemispheres.

This pattern is mechanistically interpretable and expected under a correctly functioning residualisation. Residual BERT features—by construction orthogonal to the visual feature subspace—carry semantic information that anti-predicts V1v, because the voxels that best respond to visual structure in early cortex are precisely the voxels whose residual BERT prediction is zero or negative. The magnitude of this negative effect (*g >* 6 bilaterally) confirms that the wipe procedure is operating correctly: it has removed the shared visual-semantic subspace and left a residual that is orthogonal to what early visual cortex encodes. This result functions as an internal negative control that strengthens confidence in the positive findings in EBA and lateral cortex.

(Figure 4 subject-level replication plot—EBA RH, lateral RH, V1v LH—individual subject dots with group mean, dashed zero line. All 8 subjects positive for EBA and lateral; all 8 negative for V1v.)

**Figure 4.**
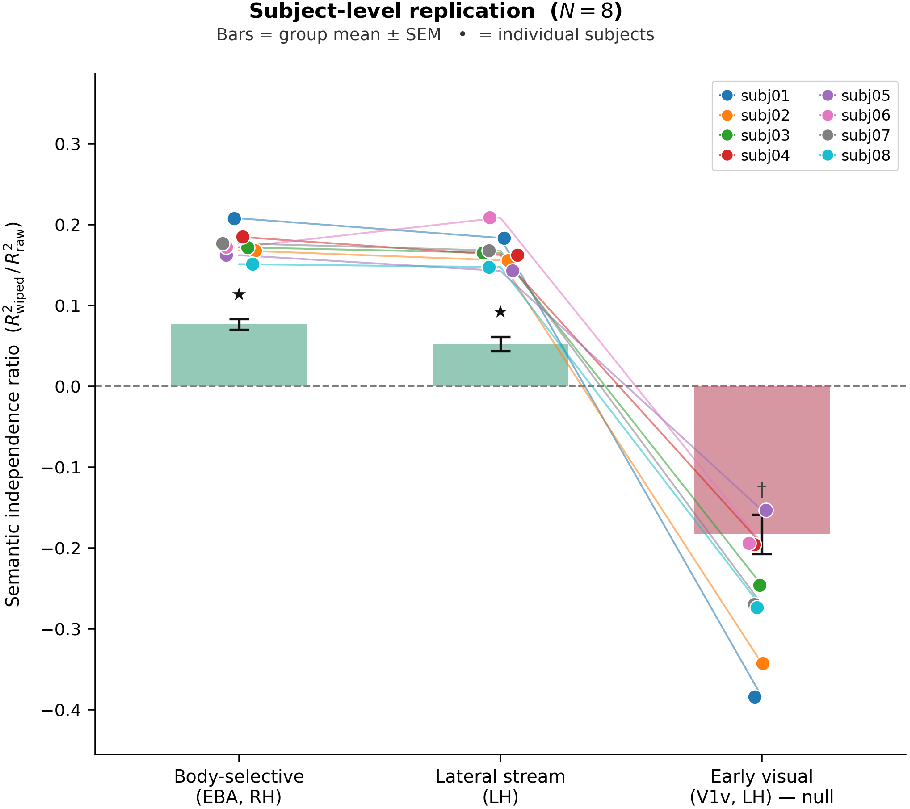
Subject-level replication of the lateral–early dissociation (*N* = 8). Semantic independence ratio 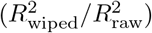 for three representative ROIs. Bars = group mean *±* SEM; dots = individual subjects (colour-coded). Body-selective cortex (EBA, RH) and lateral stream (LH) are positive for all 8 subjects; early visual cortex (V1v, LH) is negative for all 8 subjects, confirming the regional dissociation at the individual level. ⋆ = significant by both BH-FDR *t*-test and sign-flip permutation test (2^8^ permutations, *q <* 0.05); *†* = *t*-test FDR only.

### 3.5. The regional dissociation holds across multiple language models and visual feature architectures

The analyses above used BERT as the language model and a Gabor+VGG19 visual feature set. To assess whether the regional dissociation depends on these specific architectural choices, we repeated the full pipeline across a 3 × 2 factorial of language models (BERT-768, GPT-2-768, CLIP-text-512) and visual feature variants (*X*_*a*_: broad ensemble of Gabor + VGG19 + ResNet50 + CLIP-visual + EfficientNet-B0 totalling 4,352 dimensions; *X*_*b*_: VGG19 hierarchical multi-layer GAP at relu3/relu4/relu5, totalling 1,280 dimensions), yielding 6 independent wipe conditions per subject per ROI.A further validation of the residualisation framework emerges from the ordering of residual R^2^wiped values across language models. Across all ROIs, GPT-2 consistently produced the largest residual variance, followed by BERT, with CLIP-text yielding the smallest — precisely the order predicted by each model’s degree of visual grounding. GPT-2, trained exclusively on text with no visual signal, retains the most semantic content orthogonal to vision by default. BERT, trained on text but exposed indirectly to visually-grounded language, occupies an intermediate position. CLIP-text, whose text encoder is trained via direct image-text contrastive alignment to maximally overlap with visual representations, predictably yields the smallest residual after visual partialling. This theoretically motivated ordering — reproduced across both hemispheres and all ROIs — constitutes an independent validation of the pipeline: the residualisation is selectively removing visually-grounded semantic content in proportion to each model’s visual entanglement, rather than removing signal arbitrarily. Critically, despite producing the smallest absolute residuals, CLIP-text still shows positive R^2^wiped in EBA and lateral occipitotemporal cortex bilaterally, meaning the lateral-over-ventral dissociation holds even in the most conservative language model condition. If any residual semantic signal were to survive visual partialling, the CLIP-text condition is where it would be hardest to find — and it survives regardless.

The heatmap in Figure 5 displays group-mean 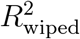 across all ROIs and all 6 wipe conditions for both hemispheres. Three results are consistent across all 6 conditions. First, EBA and lateral cortex show positive 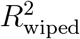 (warm colours) bilaterally in all 6 conditions, with significance markers at the strongest conditions. Second, ventral stream and place-selective regions show smaller positive effects, with the lateral *>* ventral ordering preserved in all 6 conditions. Third, early visual cortex (V1v, V1d, V2v, V2d, V3v, V3d, early stream) shows uniformly negative 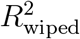 (cool colours) in all 6 conditions across both hemispheres without exception. The *X*_*a*_ conditions produce slightly larger absolute 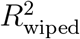 magnitudes than *X*_*b*_ conditions, as expected—a broader visual feature ensemble removes more visual variance from the language embedding, leaving a cleaner but smaller residual. The rank ordering of ROIs is fully preserved between *X*_*a*_ and *X*_*b*_ for all three language models.(See Figure 5)

**Figure 5.**
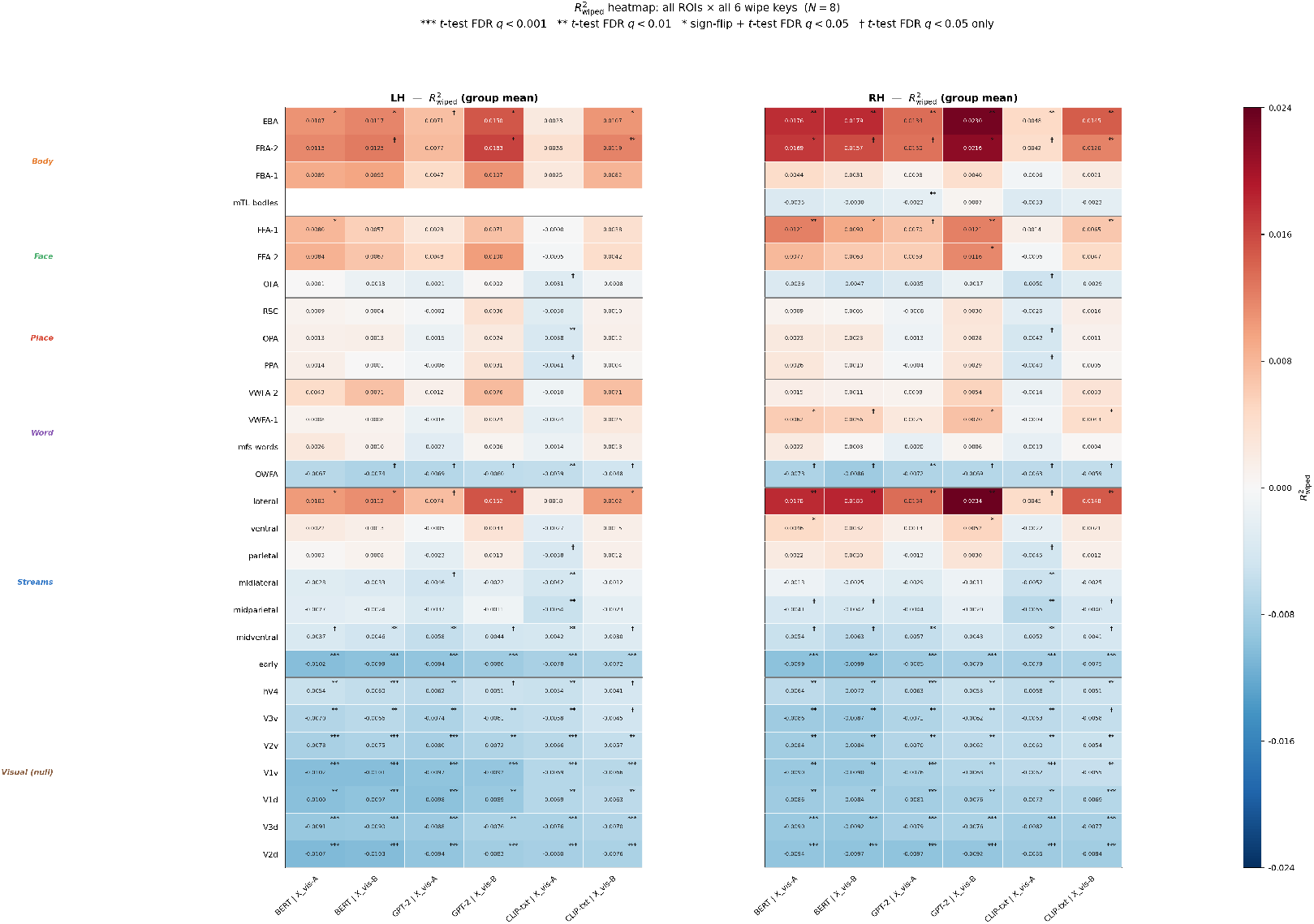
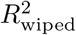 heatmap across all ROIs and wipe conditions (*N* = 8). Colour encodes the group-mean cross-validated *R*^2^ of voxelwise ridge regression predicting BOLD responses from vision-residualised language embeddings 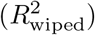, shown separately for left (LH) and right (RH) hemisphere. Rows are 28 cortical ROIs grouped by functional category (body-selective, face-selective, place-selective, word-selective, streams, retinotopic), sorted by descending 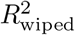 within each group. Columns correspond to the six wipe conditions: BERT, GPT-2, and CLIP-text, each residualised against two visual feature sets (*X*_vis-A_: Gabor + VGG19 + ResNet50 + CLIP-visual + EfficientNet-B0; *X*_vis-B_: Gabor + VGG19 multi-layer GAP). Cell values are group means; the diverging colour scale is centred at zero (red = positive, blue = negative). Statistical markers indicate BH-FDR corrected significance: ^∗^ sign-flip *q <* 0.05 (2^8^ permutations, requires both BH-FDR *t*-test and sign-flip); ^∗∗^ *t*-test *q <* 0.01; ^∗∗∗^ *t*-test *q <* 0.001. Body-selective (EBA, FBA-2) and lateral stream ROIs show the largest and most consistent positive values across all wipe conditions bilaterally (peak 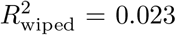, GPT-2 *X*_vis-B_, lateral RH), with 31 cells surviving sign-flip correction. Retinotopic areas (V1–V4, hV4) are uniformly negative across all 168 cells (range −0.004 to −0.011), confirming that residualised language embeddings carry no predictive signal in early visual cortex.Note: FBA-1 (*n* = 5–6), FBA-2 LH, FFA-2 LH (*n* = 7), VWFA-2 RH (*n* = 5), and mTL-bodies RH (*n* = 2) have reduced sample sizes due to ROI availability across subjects; mTL-bodies RH results should be interpreted with caution.

Across language models, even CLIP-text under *X*_*a*_ shows the largest absolute 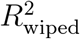 values in EBA and lateral cortex (EBA RH notably starred in the heatmap), which is despite CLIP-text being trained with direct image-text alignment and thus carrying image-predictive semantic content beyond BERT or GPT-2. Crucially, GPT-2—which has no visual training signal whatsoever—produces the strongest regional dissociation: EBA and lateral positive, ventral smaller, early visual negative. This rules out any explanation based on visually-grounded training data alone. The architecture-invariance of the regional dissociation across all 6 wipe conditions constitutes convergent validity that substantially strengthens the primary finding.

(See Figure 5)

## 4. Discussion

### 4.1. Summary of principal findings

We asked whether voxelwise responses in human visual cortex carry semantic information orthogonal to their visual feature content. Applying a cross-validated residualisation pipeline to 7T fMRI data from the Natural Scenes Dataset (Allen et al., 2022), we show the answer is yes. After removing variance in BERT representations predictable from a broad visual feature ensemble, the residual signal explains significant variance across 40 of 54 tested ROIs. Effects are largest in lateral occipitotemporal cortex, particularly EBA and the lateral stream as a whole, with results reaching the minimum achievable sign-flip *p*-value across all eight participants. Early visual cortex shows significantly negative residual prediction. This double dissociation is preserved across all six wipe conditions in a 3 × 2 factorial of language models (BERT, GPT-2, CLIP-text) and visual feature architectures, establishing architecture-invariance. The principal methodological contribution is an approach that explicitly separates language-model variance from visual-feature variance in voxelwise encoding models—a dissociation that prior alignment studies (Doerig et al., 2025; Marcos-Manchón and Fuentemilla, 2025) were not designed to make.

### 4.2. Why lateral occipitotemporal cortex carries non-visual semantic information

EBA is canonically characterised as body-selective by contrast with objects, faces, and scenes (Downing et al., 2001), so the magnitude of the non-visual residual there is not self-evident. It is, however, interpretable given converging evidence that EBA participates in a broader semantic network. Using fMRI adaptation, (Kubiak and Króliczak, 2016) demonstrated that left EBA tracks gesture semantics rather than visual similarity. (Marrazzo et al., 2023) showed that purely visual models leave substantial unexplained variance in EBA encoding of body stimuli. Most directly, (Gandolfo et al., 2024) showed using a large dataset (*n* = 92) and fMRI-guided TMS that left EBA responds selectively to face-to-face dyads and that stimulation causally abolishes efficient perception of social interactions—establishing EBA as a locus of relational body processing. The non-visual language-model variance we observe is consistent with EBA encoding thematic and social-relational content that is not recoverable from image content by the visual models tested; whether a more expressive visual architecture could account for this variance remains an open question. This interpretation is further supported by (Wurm et al., 2017), who showed that lateral regions represent action-related and animate semantic categories, and by (Contier et al., 2024), whose 66 behaviour-derived dimensions implicate EBA and face-selective regions in social and agentive tuning. Our contribution is to demonstrate that a portion of this tuning—approximately 17–19% of total explainable variance in LOTC—is orthogonal to visual features, a claim not previously testable without explicit visual partialling.

### 4.3. The negative V1v result as mechanistic evidence

V1v shows significantly negative residual prediction across all eight subjects, with effect sizes substantially larger than the positive effects in EBA. This is mechanistically expected: the visual feature space was optimised to capture visual content in BERT embeddings, so the residual lives in a subspace geometrically orthogonal to—and therefore negatively correlated with—what V1v responds to. Critically, this provides an internal validity check. Miscalibrated residualisation would produce broadly positive residuals including in early visual cortex. The strongly negative V1 result, with virtually no voxel showing positive residual prediction in any subject, demonstrates that the residual is genuinely post-visual rather than a contaminated signal. The sharp contrast with the predominantly positive residuals in EBA underscores that the regional pattern reflects a real structural property of the data.

### 4.4. Relation to prior work on language model alignment with visual cortex

(Doerig et al., 2025) showed that LLM embeddings of scene captions predict activity throughout visual cortex, capturing area-specific selectivities. (Marcos-Manchón and Fuentemilla, 2025) characterised lateral and ventral scene-processing routes using inter-subject RSA, finding language-model alignment concentrated in lateral LOTC. Our spatial localisation is consistent with both. The critical distinction is methodological: standard RSA alignment is ambiguous when language and vision models share representations of the same scenes. By explicitly partialling visual-feature variance, we show that residual language-model variance—orthogonal to what visual models explain—remains significant in EBA and lateral LOTC. That GPT-2, which has no visual training signal whatsoever, produces the same regional dissociation as BERT and CLIP-text rules out explanations based on shared visual training data and points instead to something about the semantic content that LOTC represents. (Contier et al., 2024) establish that semantic tuning is present broadly in visual cortex; the present approach establishes that in lateral LOTC this tuning is partly independent of visual features.

### 4.5. Candidate mechanisms

The residual language-model signal in EBA could reflect several non-mutually-exclusive properties. First, **relational and compositional structure**: visual models capture local feature statistics but struggle to encode relational propositions — a nurse chasing a man and a man chasing a nurse have similar pixel statistics, but language models encode the relational structure with genuine sensitivity (Abassi and Papeo, 2024, 2020). Because EBA has been shown to represent two-body configurations and dyadic social interactions, its residual may reflect this relational content. Second, **social role and thematic context**: a nurse and a patient may be visually indistinguishable but have distinct language-model representations; body configurations proposed to encode dyadic social roles (Gandolfo et al., 2024) would produce exactly this pattern after visual partialling. The concentration of effects in lateral rather than ventral cortex—where scene-context and geometry dimensions dominate—favours relational and social-role accounts over purely affective (de Gelder, 2015) or thematic explanations.

### 4.6. VWFA-2 secondary finding

Left VWFA-2 shows significant non-visual language-model variance, confirmed by sign-flip test across all eight subjects (*q*_signflip_ = 0.020), with a semantic independence ratio of 0.172 comparable to EBA. The right-hemisphere analogue did not survive sign-flip correction (*N* = 5, *q*_signflip_ = 0.065) and is not discussed further. VWFA-2 has documented sensitivity to lexical and semantic properties beyond visual word form such as word and letter string statistics (Vinckier et al., 2007; Woolnough et al., 2021; Kronbichler et al., 2004). That this extends to natural scene images is consistent with its responsiveness to auditory language, suggesting it may not be strictly, though primarily visual (Li et al., 2024). Alternatively, NSD images containing visible text — books, signs, screens — may drive spurious visually-independent predictions in a region tuned to orthographic features. Disambiguating these accounts would require restricting the analysis to trials without text-bearing objects, which we leave for future work. We do not interpret this observation further here.

### 4.7. Limitations

Three limitations warrant emphasis. First, this study cannot establish the directionality of the visual-to-semantic interface. Post-visual language-model variance is compatible with bottom-up semantic inference, top-down feedback from language or memory networks, or intrinsic representational geometry reflecting the region’s position at the border of visual and semantic cortex—all three are consistent with our results. Second, shared training data biases cannot be fully excluded: BERT corpora and MS-COCO captions both describe natural scenes. The strongly negative V1 result argues against a uniform bias as the dominant explanation, but a partial contribution cannot be ruled out (Doerig et al., 2025). Third, *n* = 8 limits sign-flip resolution to *p*_min_ = 0.0039, and a single train-test split was used for computational tractability; neither materially affects conclusions for the principal findings given the observed effect sizes.

A common question when interpreting the results of variance partitioning is whether the remaining variance in the language model reflects true amodal semantic content, feedback from language networks in the frontal or temporal regions, or some intrinsic geometry of a region at the boundary of the visual and semantic cortex. We choose not to settle this debate because that would involve addressing a deeper philosophical question about where visual information ends and semantic information starts. The current data cannot answer that, and it remains a topic of discussion in the broader literature. Our approach avoids this boundary issue by design: we measure how separate the language model embeddings are from a specific visual feature subspace and explore whether this separate component predicts neural responses. This perspective does not take a stand on the type of representation involved; it makes no assertion about whether the remaining signal is genuinely amodal, nor does it require visual and semantic representations to be completely different. We see this uncertainty as a strength rather than a weakness. Since the finding focuses on measurable separation instead of a theoretical position, it is resilient to different explanations of the visual-semantic relationship and can be understood within various theoretical frameworks as they develop.

### 4.8. Broader implications

The dominant framework treats the semantic tuning of higher visual areas as a consequence of their visual selectivity (Contier et al., 2024). Our results are compatible with this but require an extension: a non-trivial fraction of language-model variance in lateral LOTC persists after controlling for visual features, suggesting these regions may represent information accessible from multiple modalities (Huth et al., 2016). For visual encoding models, the implication is practical: feedforward visual feature models systematically underpredict the language-aligned content of body-selective and social-perception regions. Incorporating large language model embeddings alongside visual features (Doerig et al., 2025)—and, importantly, using the residualisation approach introduced here to isolate their independent contributions— may substantially improve prediction ceilings in lateral LOTC.

## Data and Code Availability

The fMRI data used in this study are from the Natural Scenes Dataset (NSD), a publicly available resource accessible at http://naturalscenesdataset.org Allen et al. (2022). No new data were collected or generated for this study. The MS-COCO image annotations and captions used as semantic descriptors are available at https://cocodataset.org.

All analysis code, including the visual residualisation pipeline, voxelwise encoding models, ROI aggregation, and statistical procedures, is publicly available at. The repository includes scripts for reproducing all the statistical results reported in this paper.

## Declaration of Generative AI and AI-Assisted Technologies in the Manuscript Preparation Process

During the preparation of this work, the author(s) used **Claude Sonnet 4.6 (Anthropic)** in order to assist with *articulation, writing coherent sentences, and conversion of content to L*^*A*^*TEX code*. After using this tool/service, the author(s) reviewed and edited the content as needed and take(s) full responsibility for the content of the published article.

## Supplementary Material

The supplementary material for this article contains extended data tables and robustness analyses supporting the main findings.

*Supplementary Table 1: Subject-Level Visually-Independent Semantic Encoding Across Visual ROIs*

Table S1 reports subject-level estimates of visually-independent semantic encoding 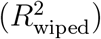 across all visual regions of interest (ROIs) for each of the eight participants. For every subject–hemisphere–ROI combination, we provide (i) the number of voxels (*N*_vox_), (ii) the mean *R*^2^ after visual variance removal 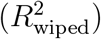, (iii) the raw encoding strength 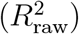, (iv) their ratio 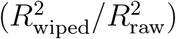, and (v) group-level significance flags derived from a one-sample *t*-test and a sign-flip permutation test, both corrected for multiple comparisons using the Benjamini–Hochberg false-discovery rate (BH-FDR) procedure.

ROIs are grouped into the following functional categories: stream-level regions (early, lateral, mid-lateral, mid-parietal, mid-ventral, parietal, and ventral streams), body-selective regions (EBA, FBA-1, FBA-2, mTL-bodies), face-selective regions (FFA-1, FFA-2, OFA), place-selective regions (OPA, PPA, RSC), word-selective regions (OWFA, VWFA-1, VWFA-2, mTL-words, mfs-words), and early retinotopic visual areas (V1d, V1v, V2d, V2v, V3d, V3v, hV4). Consistent with the main text, positive and significant 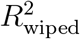 values are concentrated in lateral-stream, body-selective, and face-selective ROIs, while early visual areas exhibit uniformly negative values, confirming that residualised variance in those regions reflects suppression rather than semantic encoding.

*Supplementary Table 2: Group-Level Visually-Independent Semantic Encoding Across Models and ROIs*

Table S2 extends the subject-level results by reporting group-level summaries (mean *±* SEM across subjects) of 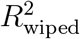 for all combinations of ROI, hemisphere, and language/vision model variant. Three language models are compared—BERT, GPT-2, and CLIP-text—each evaluated under two visual residualisation strategies: *X*_vis_-A (a broad ensemble comprising Gabor filters, VGG19, ResNet50, CLIP-visual, and EfficientNet-B0) and *X*_vis_-B (a VGG19 hierarchical multi-layer global average pooling approach). In addition to mean 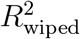 and the ratio to 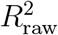, the table provides Hedges’ *g* as a measure of effect size and BH-FDR-corrected *p*-values from both the one-sample *t*-test (*q*_FDR_) and the sign-flip permutation test (*q*_SF_).

*Supplementary Figure S1: Effect of Visual Residualisation Strategy on* 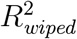

Figure S1 assesses whether the choice of visual feature set affects the rank ordering of ROIs in terms of visually-independent semantic encoding. Each panel plots 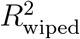 obtained under *X*_vis_-A residualisation against the corresponding values under *X*_vis_-B, separately for BERT, GPT-2, and CLIP-text across both hemispheres (*N* = 8 per panel). Individual ROIs are colour-coded by functional group. Pearson correlations between the two residualisation conditions are uniformly high (*r* = 0.977–0.998, all *p <* 10^−17^), confirming that the ROI rank ordering is robust to the choice of visual feature set. GPT-2 and CLIP-text show a systematic above-diagonal shift (*X*_vis_-B *> X*_vis_-A in 27–29 of 27–29 ROIs per hemisphere), while BERT displays no consistent directional preference. Deviations from the identity line are small in all cases (mean | *X*_vis_-B − *X*_vis_-A | *<* 0.001), supporting the equivalence of both strategies for the purposes of the main analysis.

