## Supplemental Material for "Semantic Information Orthogonal to Visual Features Peaks in Lateral Occipitotemporal Cortex"

Arun Ram Ponnambalam

*Department of Biomedical Engineering, SRM Institute of Science and Technology (SRMIST), , Chengalpattu, , Tamil Nadu, India*

Krishnan Pottore Venkiteswaran

*Department of Computer Science, University College Dublin, , Dublin, Ireland*

*Keywords:*

### Supplementary Material

Table 1: **Subject-level visually-independent semantic encoding ($R^2_{\text{wiped}}$) across visual ROIs.** Each row is one subject × hemisphere × ROI observation. $R^2_{\text{wiped}}$: subject-level mean $R^2$ after visual variance removal. $R^2_{\text{raw}}$: subject-level mean raw $R^2$. Ratio = $R^2_{\text{wiped}}/R^2_{\text{raw}}$. Sig. $t$-test and Sig. SF reflect group-level significance flags. Rows sorted alphabetically by ROI then hemisphere then subject within each category.

| **ROI** | **H** | **Subject** | $N_{\text{vox}}$ | $R^2_{\text{wiped}}$ | $R^2_{\text{raw}}$ | **Ratio** | **Sig. $t$-test** | **Sig. SF** |
|---|---|---|---|---|---|---|---|---|
| *Stream-level ROIs* | | | | | | | | |
| Early stream | LH | subj01 | 5169 | −0.0132 | 0.040 | −0.331 | True | False |
| Early stream | LH | subj02 | 5171 | −0.0130 | 0.047 | −0.279 | True | False |
| Early stream | LH | subj03 | 5168 | −0.0107 | 0.043 | −0.247 | True | False |
| Early stream | LH | subj04 | 5170 | −0.0109 | 0.042 | −0.261 | True | False |
| Early stream | LH | subj05 | 5169 | −0.0083 | 0.088 | −0.095 | True | False |
| Early stream | LH | subj06 | 5141 | −0.0079 | 0.052 | −0.154 | True | False |
| Early stream | LH | subj07 | 5169 | −0.0104 | 0.037 | −0.279 | True | False |

*Continued on next page*

Table 1 – *continued from previous page*

| **ROI** | **H** | **Subject** | $N_{\text{vox}}$ | $R^2_{\text{wiped}}$ | $R^2_{\text{raw}}$ | **Ratio** | **Sig. $t$-test** | **Sig. SF** |
|---|---|---|---|---|---|---|---|---|
| Early stream | LH | subj08 | 5146 | −0.0110 | 0.039 | −0.280 | True | False |
| Early stream | RH | subj01 | 5304 | −0.0130 | 0.039 | −0.338 | True | False |
| Early stream | RH | subj02 | 5300 | −0.0133 | 0.048 | −0.276 | True | False |
| Early stream | RH | subj03 | 5302 | −0.0097 | 0.037 | −0.264 | True | False |
| Early stream | RH | subj04 | 5304 | −0.0101 | 0.045 | −0.223 | True | False |
| Early stream | RH | subj05 | 5302 | −0.0076 | 0.082 | −0.093 | True | False |
| Early stream | RH | subj06 | 4978 | −0.0086 | 0.058 | −0.149 | True | False |
| Early stream | RH | subj07 | 5302 | −0.0099 | 0.036 | −0.274 | True | False |
| Early stream | RH | subj08 | 5286 | −0.0111 | 0.035 | −0.317 | True | False |
| Lateral stream | LH | subj01 | 3129 | 0.0442 | 0.241 | 0.184 | True | True |
| Lateral stream | LH | subj02 | 3127 | 0.0326 | 0.209 | 0.156 | True | True |
| Lateral stream | LH | subj03 | 3128 | 0.0250 | 0.152 | 0.165 | True | True |
| Lateral stream | LH | subj04 | 3126 | 0.0212 | 0.130 | 0.162 | True | True |
| Lateral stream | LH | subj05 | 3136 | 0.0369 | 0.258 | 0.143 | True | True |
| Lateral stream | LH | subj06 | 3128 | 0.0282 | 0.135 | 0.209 | True | True |
| Lateral stream | LH | subj07 | 3124 | 0.0318 | 0.189 | 0.168 | True | True |
| Lateral stream | LH | subj08 | 3129 | 0.0123 | 0.084 | 0.148 | True | True |
| Lateral stream | RH | subj01 | 3578 | 0.0518 | 0.244 | 0.212 | True | True |
| Lateral stream | RH | subj02 | 3573 | 0.0476 | 0.280 | 0.170 | True | True |
| Lateral stream | RH | subj03 | 3577 | 0.0336 | 0.190 | 0.177 | True | True |
| Lateral stream | RH | subj04 | 3576 | 0.0544 | 0.267 | 0.204 | True | True |
| Lateral stream | RH | subj05 | 3574 | 0.0533 | 0.292 | 0.183 | True | True |
| Lateral stream | RH | subj06 | 3577 | 0.0327 | 0.183 | 0.179 | True | True |
| Lateral stream | RH | subj07 | 3574 | 0.0345 | 0.201 | 0.172 | True | True |
| Lateral stream | RH | subj08 | 3578 | 0.0153 | 0.099 | 0.154 | True | True |
| Mid-lateral | LH | subj01 | 1091 | 0.0017 | 0.076 | 0.023 | False | False |
| Mid-lateral | LH | subj02 | 1100 | −0.0055 | 0.091 | −0.060 | False | False |
| Mid-lateral | LH | subj03 | 1095 | 0.0004 | 0.097 | 0.004 | False | False |
| Mid-lateral | LH | subj04 | 1096 | 0.0009 | 0.083 | 0.011 | False | False |
| Mid-lateral | LH | subj05 | 1098 | 0.0107 | 0.136 | 0.079 | False | False |
| Mid-lateral | LH | subj06 | 1097 | 0.0169 | 0.129 | 0.131 | False | False |
| Mid-lateral | LH | subj07 | 1098 | 0.0011 | 0.062 | 0.018 | False | False |
| Mid-lateral | LH | subj08 | 1093 | 0.0020 | 0.078 | 0.025 | False | False |
| Mid-lateral | RH | subj01 | 1191 | 0.0045 | 0.104 | 0.043 | False | False |

*Continued on next page*

Table 1 – *continued from previous page*

| **ROI** | **H** | **Subject** | $N_{\text{vox}}$ | $R^2_{\text{wiped}}$ | $R^2_{\text{raw}}$ | **Ratio** | **Sig.** $t$**-test** | **Sig. SF** |
|---|---|---|---|---|---|---|---|---|
| Mid-lateral | RH | subj02 | 1194 | $-0.0040$ | 0.082 | $-0.048$ | False | False |
| Mid-lateral | RH | subj03 | 1192 | 0.0012 | 0.068 | 0.018 | False | False |
| Mid-lateral | RH | subj04 | 1190 | 0.0069 | 0.121 | 0.057 | False | False |
| Mid-lateral | RH | subj05 | 1188 | 0.0193 | 0.189 | 0.102 | False | False |
| Mid-lateral | RH | subj06 | 1190 | 0.0122 | 0.131 | 0.094 | False | False |
| Mid-lateral | RH | subj07 | 1190 | 0.0000 | 0.085 | 0.000 | False | False |
| Mid-lateral | RH | subj08 | 1188 | $-0.0036$ | 0.041 | $-0.088$ | False | False |
| Mid-parietal | LH | subj01 | 853 | 0.0011 | 0.117 | 0.009 | False | False |
| Mid-parietal | LH | subj02 | 852 | $-0.0067$ | 0.072 | $-0.093$ | False | False |
| Mid-parietal | LH | subj03 | 852 | $-0.0016$ | 0.065 | $-0.024$ | False | False |
| Mid-parietal | LH | subj04 | 853 | $-0.0026$ | 0.082 | $-0.032$ | False | False |
| Mid-parietal | LH | subj05 | 854 | 0.0187 | 0.231 | 0.081 | False | False |
| Mid-parietal | LH | subj06 | 853 | $-0.0015$ | 0.064 | $-0.023$ | False | False |
| Mid-parietal | LH | subj07 | 857 | $-0.0010$ | 0.082 | $-0.012$ | False | False |
| Mid-parietal | LH | subj08 | 851 | $-0.0008$ | 0.069 | $-0.012$ | False | False |
| Mid-parietal | RH | subj01 | 1133 | 0.0022 | 0.100 | 0.022 | False | False |
| Mid-parietal | RH | subj02 | 1131 | $-0.0061$ | 0.076 | $-0.079$ | False | False |
| Mid-parietal | RH | subj03 | 1134 | $-0.0044$ | 0.064 | $-0.068$ | False | False |
| Mid-parietal | RH | subj04 | 1130 | $-0.0020$ | 0.081 | $-0.025$ | False | False |
| Mid-parietal | RH | subj05 | 1138 | 0.0093 | 0.173 | 0.053 | False | False |
| Mid-parietal | RH | subj06 | 1134 | 0.0009 | 0.083 | 0.011 | False | False |
| Mid-parietal | RH | subj07 | 1132 | 0.0044 | 0.048 | 0.093 | False | False |
| Mid-parietal | RH | subj08 | 1133 | $-0.0051$ | 0.054 | $-0.093$ | False | False |
| Mid-ventral | LH | subj01 | 867 | $-0.0064$ | 0.070 | $-0.092$ | False | False |
| Mid-ventral | LH | subj02 | 862 | $-0.0080$ | 0.079 | $-0.101$ | False | False |
| Mid-ventral | LH | subj03 | 867 | $-0.0018$ | 0.079 | $-0.023$ | False | False |
| Mid-ventral | LH | subj04 | 868 | $-0.0038$ | 0.069 | $-0.055$ | False | False |
| Mid-ventral | LH | subj05 | 863 | 0.0073 | 0.152 | 0.048 | False | False |
| Mid-ventral | LH | subj06 | 868 | 0.0106 | 0.128 | 0.083 | False | False |
| Mid-ventral | LH | subj07 | 867 | $-0.0056$ | 0.062 | $-0.092$ | False | False |
| Mid-ventral | LH | subj08 | 862 | $-0.0078$ | 0.064 | $-0.122$ | False | False |
| Mid-ventral | RH | subj01 | 1050 | $-0.0072$ | 0.049 | $-0.147$ | False | False |
| Mid-ventral | RH | subj02 | 1051 | $-0.0089$ | 0.055 | $-0.160$ | False | False |
| Mid-ventral | RH | subj03 | 1049 | 0.0004 | 0.081 | 0.006 | False | False |

*Continued on next page*

Table 1 – *continued from previous page*

| **ROI** | **H** | **Subject** | $N_{\text{vox}}$ | $R^2_{\text{wiped}}$ | $R^2_{\text{raw}}$ | **Ratio** | **Sig.** $t$**-test** | **Sig. SF** |
|---|---|---|---|---|---|---|---|---|
| Mid-ventral | RH | subj04 | 1050 | 0.0001 | 0.083 | 0.001 | False | False |
| Mid-ventral | RH | subj05 | 1050 | 0.0158 | 0.191 | 0.083 | False | False |
| Mid-ventral | RH | subj06 | 1049 | 0.0039 | 0.088 | 0.044 | False | False |
| Mid-ventral | RH | subj07 | 1049 | −0.0048 | 0.058 | −0.083 | False | False |
| Mid-ventral | RH | subj08 | 1049 | −0.0055 | 0.061 | −0.091 | False | False |
| Parietal stream | LH | subj01 | 2346 | 0.0159 | 0.142 | 0.112 | True | True |
| Parietal stream | LH | subj02 | 2343 | 0.0070 | 0.112 | 0.063 | True | True |
| Parietal stream | LH | subj03 | 2345 | 0.0068 | 0.100 | 0.068 | True | True |
| Parietal stream | LH | subj04 | 2344 | 0.0105 | 0.084 | 0.124 | True | True |
| Parietal stream | LH | subj05 | 2336 | 0.0179 | 0.176 | 0.102 | True | True |
| Parietal stream | LH | subj06 | 2343 | 0.0114 | 0.088 | 0.130 | True | True |
| Parietal stream | LH | subj07 | 2340 | 0.0098 | 0.090 | 0.109 | True | True |
| Parietal stream | LH | subj08 | 2345 | −0.0006 | 0.045 | −0.013 | True | True |
| Parietal stream | RH | subj01 | 2314 | 0.0166 | 0.122 | 0.136 | True | True |
| Parietal stream | RH | subj02 | 2319 | 0.0083 | 0.125 | 0.067 | True | True |
| Parietal stream | RH | subj03 | 2315 | 0.0053 | 0.087 | 0.061 | True | True |
| Parietal stream | RH | subj04 | 2318 | 0.0099 | 0.077 | 0.129 | True | True |
| Parietal stream | RH | subj05 | 2318 | 0.0256 | 0.203 | 0.126 | True | True |
| Parietal stream | RH | subj06 | 2316 | 0.0091 | 0.075 | 0.122 | True | True |
| Parietal stream | RH | subj07 | 2319 | 0.0190 | 0.088 | 0.215 | True | True |
| Parietal stream | RH | subj08 | 2317 | 0.0022 | 0.046 | 0.049 | True | True |
| Ventral stream | LH | subj01 | 4974 | 0.0137 | 0.150 | 0.091 | True | True |
| Ventral stream | LH | subj02 | 4974 | 0.0133 | 0.216 | 0.062 | True | True |
| Ventral stream | LH | subj03 | 4975 | 0.0104 | 0.128 | 0.081 | True | True |
| Ventral stream | LH | subj04 | 4972 | 0.0119 | 0.136 | 0.087 | True | True |
| Ventral stream | LH | subj05 | 4972 | 0.0284 | 0.254 | 0.112 | True | True |
| Ventral stream | LH | subj06 | 4973 | 0.0112 | 0.104 | 0.107 | True | True |
| Ventral stream | LH | subj07 | 4972 | 0.0066 | 0.116 | 0.057 | True | True |
| Ventral stream | LH | subj08 | 4980 | 0.0043 | 0.085 | 0.050 | True | True |
| Ventral stream | RH | subj01 | 4852 | 0.0155 | 0.180 | 0.086 | True | True |
| Ventral stream | RH | subj02 | 4854 | 0.0090 | 0.180 | 0.050 | True | True |
| Ventral stream | RH | subj03 | 4853 | 0.0162 | 0.164 | 0.099 | True | True |
| Ventral stream | RH | subj04 | 4855 | 0.0125 | 0.155 | 0.081 | True | True |
| Ventral stream | RH | subj05 | 4852 | 0.0224 | 0.218 | 0.103 | True | True |

*Continued on next page*

Table 1 – *continued from previous page*

| **ROI** | **H** | **Subject** | $N_{\text{vox}}$ | $R^2_{\text{wiped}}$ | $R^2_{\text{raw}}$ | **Ratio** | **Sig.** $t$**-test** | **Sig. SF** |
|---|---|---|---|---|---|---|---|---|
| Ventral stream | RH | subj06 | 4852 | 0.0131 | 0.140 | 0.093 | True | True |
| Ventral stream | RH | subj07 | 4852 | 0.0115 | 0.137 | 0.084 | True | True |
| Ventral stream | RH | subj08 | 4854 | 0.0100 | 0.114 | 0.087 | True | True |
| *Body-selective* | | | | | | | | |
| EBA | LH | subj01 | 2837 | 0.0473 | 0.247 | 0.191 | True | True |
| EBA | LH | subj02 | 3059 | 0.0310 | 0.209 | 0.149 | True | True |
| EBA | LH | subj03 | 3017 | 0.0241 | 0.160 | 0.151 | True | True |
| EBA | LH | subj04 | 3538 | 0.0202 | 0.140 | 0.144 | True | True |
| EBA | LH | subj05 | 3618 | 0.0344 | 0.245 | 0.141 | True | True |
| EBA | LH | subj06 | 3605 | 0.0300 | 0.151 | 0.199 | True | True |
| EBA | LH | subj07 | 3873 | 0.0261 | 0.168 | 0.156 | True | True |
| EBA | LH | subj08 | 3349 | 0.0122 | 0.090 | 0.135 | True | True |
| EBA | RH | subj01 | 3400 | 0.0550 | 0.265 | 0.208 | True | True |
| EBA | RH | subj02 | 3650 | 0.0479 | 0.285 | 0.168 | True | True |
| EBA | RH | subj03 | 3568 | 0.0328 | 0.191 | 0.172 | True | True |
| EBA | RH | subj04 | 4562 | 0.0463 | 0.251 | 0.185 | True | True |
| EBA | RH | subj05 | 5741 | 0.0431 | 0.266 | 0.162 | True | True |
| EBA | RH | subj06 | 4060 | 0.0349 | 0.203 | 0.172 | True | True |
| EBA | RH | subj07 | 3381 | 0.0380 | 0.215 | 0.177 | True | True |
| EBA | RH | subj08 | 3425 | 0.0156 | 0.103 | 0.151 | True | True |
| FBA-1 | LH | subj01 | 574 | 0.0325 | 0.192 | 0.170 | True | False |
| FBA-1 | LH | subj03 | 524 | 0.0298 | 0.132 | 0.226 | True | False |
| FBA-1 | LH | subj05 | 512 | 0.0135 | 0.165 | 0.082 | True | False |
| FBA-1 | LH | subj06 | 615 | 0.0374 | 0.191 | 0.196 | True | False |
| FBA-1 | LH | subj08 | 188 | 0.0181 | 0.075 | 0.242 | True | False |
| FBA-1 | RH | subj01 | 206 | −0.0036 | 0.099 | −0.036 | True | False |
| FBA-1 | RH | subj03 | 1176 | 0.0164 | 0.132 | 0.124 | True | False |
| FBA-1 | RH | subj04 | 395 | 0.0056 | 0.138 | 0.040 | True | False |
| FBA-1 | RH | subj05 | 556 | 0.0207 | 0.200 | 0.103 | True | False |
| FBA-1 | RH | subj06 | 1065 | 0.0180 | 0.140 | 0.128 | True | False |
| FBA-1 | RH | subj08 | 923 | 0.0193 | 0.131 | 0.147 | True | False |
| FBA-2 | LH | subj02 | 1058 | 0.0287 | 0.218 | 0.132 | True | True |
| FBA-2 | LH | subj03 | 92 | 0.0526 | 0.245 | 0.214 | True | True |
| FBA-2 | LH | subj04 | 787 | 0.0226 | 0.110 | 0.204 | True | True |

*Continued on next page*

Table 1 – *continued from previous page*

| ROI | H | Subject | $N_{\text{vox}}$ | $R^2_{\text{wiped}}$ | $R^2_{\text{raw}}$ | Ratio | Sig. $t$-test | Sig. SF |
|---|---|---|---|---|---|---|---|---|
| FBA-2 | LH | subj05 | 984 | 0.0526 | 0.302 | 0.174 | True | True |
| FBA-2 | LH | subj06 | 387 | 0.0228 | 0.087 | 0.263 | True | True |
| FBA-2 | LH | subj07 | 422 | 0.0277 | 0.144 | 0.192 | True | True |
| FBA-2 | LH | subj08 | 519 | 0.0253 | 0.155 | 0.163 | True | True |
| FBA-2 | RH | subj01 | 856 | 0.0411 | 0.270 | 0.152 | True | True |
| FBA-2 | RH | subj02 | 748 | 0.0190 | 0.189 | 0.101 | True | True |
| FBA-2 | RH | subj03 | 415 | 0.0605 | 0.333 | 0.181 | True | True |
| FBA-2 | RH | subj04 | 473 | 0.0175 | 0.145 | 0.121 | True | True |
| FBA-2 | RH | subj05 | 200 | 0.0559 | 0.359 | 0.156 | True | True |
| FBA-2 | RH | subj06 | 219 | 0.0360 | 0.215 | 0.167 | True | True |
| FBA-2 | RH | subj07 | 522 | 0.0409 | 0.252 | 0.162 | True | True |
| FBA-2 | RH | subj08 | 569 | 0.0231 | 0.143 | 0.162 | True | True |
| mTL-bodies | RH | subj06 | 112 | 0.0069 | 0.065 | 0.106 | False | False |
| mTL-bodies | RH | subj08 | 164 | 0.0052 | 0.035 | 0.150 | False | False |
| *Face-selective* | | | | | | | | |
| FFA-1 | LH | subj01 | 552 | 0.0281 | 0.218 | 0.129 | True | True |
| FFA-1 | LH | subj02 | 244 | 0.0104 | 0.144 | 0.073 | True | True |
| FFA-1 | LH | subj03 | 549 | 0.0178 | 0.159 | 0.112 | True | True |
| FFA-1 | LH | subj04 | 599 | 0.0094 | 0.137 | 0.069 | True | True |
| FFA-1 | LH | subj05 | 542 | 0.0305 | 0.250 | 0.122 | True | True |
| FFA-1 | LH | subj06 | 435 | 0.0385 | 0.234 | 0.165 | True | True |
| FFA-1 | LH | subj07 | 484 | 0.0102 | 0.106 | 0.096 | True | True |
| FFA-1 | LH | subj08 | 776 | 0.0060 | 0.097 | 0.062 | True | True |
| FFA-1 | RH | subj01 | 330 | 0.0243 | 0.187 | 0.130 | True | True |
| FFA-1 | RH | subj02 | 594 | 0.0075 | 0.157 | 0.048 | True | True |
| FFA-1 | RH | subj03 | 1067 | 0.0383 | 0.237 | 0.162 | True | True |
| FFA-1 | RH | subj04 | 526 | 0.0176 | 0.172 | 0.103 | True | True |
| FFA-1 | RH | subj05 | 658 | 0.0227 | 0.202 | 0.112 | True | True |
| FFA-1 | RH | subj06 | 167 | 0.0344 | 0.208 | 0.166 | True | True |
| FFA-1 | RH | subj07 | 601 | 0.0408 | 0.262 | 0.156 | True | True |
| FFA-1 | RH | subj08 | 577 | 0.0250 | 0.185 | 0.135 | True | True |
| FFA-2 | LH | subj02 | 401 | 0.0193 | 0.185 | 0.104 | True | True |
| FFA-2 | LH | subj03 | 125 | 0.0465 | 0.207 | 0.224 | True | True |
| FFA-2 | LH | subj04 | 434 | 0.0266 | 0.143 | 0.187 | True | True |

*Continued on next page*

Table 1 – *continued from previous page*

| **ROI** | **H** | **Subject** | $N_{\text{vox}}$ | $R^2_{\text{wiped}}$ | $R^2_{\text{raw}}$ | **Ratio** | **Sig. $t$-test** | **Sig. SF** |
|---|---|---|---|---|---|---|---|---|
| FFA-2 | LH | subj05 | 335 | 0.0534 | 0.334 | 0.160 | True | True |
| FFA-2 | LH | subj06 | 320 | 0.0122 | 0.072 | 0.169 | True | True |
| FFA-2 | LH | subj07 | 39 | −0.0046 | 0.027 | −0.168 | True | True |
| FFA-2 | LH | subj08 | 678 | 0.0097 | 0.110 | 0.088 | True | True |
| FFA-2 | RH | subj01 | 629 | 0.0443 | 0.300 | 0.148 | True | True |
| FFA-2 | RH | subj02 | 527 | 0.0147 | 0.174 | 0.084 | True | True |
| FFA-2 | RH | subj03 | 11 | 0.0194 | 0.123 | 0.158 | True | True |
| FFA-2 | RH | subj04 | 467 | 0.0193 | 0.158 | 0.122 | True | True |
| FFA-2 | RH | subj05 | 650 | 0.0325 | 0.248 | 0.131 | True | True |
| FFA-2 | RH | subj06 | 523 | 0.0263 | 0.174 | 0.152 | True | True |
| FFA-2 | RH | subj07 | 18 | −0.0058 | 0.024 | −0.241 | True | True |
| FFA-2 | RH | subj08 | 914 | 0.0199 | 0.142 | 0.140 | True | True |
| OFA | LH | subj01 | 432 | −0.0032 | 0.069 | −0.046 | False | False |
| OFA | LH | subj02 | 493 | −0.0087 | 0.071 | −0.122 | False | False |
| OFA | LH | subj03 | 607 | 0.0122 | 0.128 | 0.095 | False | False |
| OFA | LH | subj04 | 623 | 0.0021 | 0.102 | 0.021 | False | False |
| OFA | LH | subj05 | 778 | 0.0194 | 0.202 | 0.096 | False | False |
| OFA | LH | subj06 | 419 | 0.0198 | 0.159 | 0.124 | False | False |
| OFA | LH | subj07 | 434 | 0.0133 | 0.151 | 0.088 | False | False |
| OFA | LH | subj08 | 452 | −0.0012 | 0.073 | −0.016 | False | False |
| OFA | RH | subj01 | 305 | −0.0006 | 0.075 | −0.008 | False | False |
| OFA | RH | subj02 | 795 | −0.0062 | 0.046 | −0.137 | False | False |
| OFA | RH | subj03 | 836 | 0.0067 | 0.115 | 0.058 | False | False |
| OFA | RH | subj04 | 774 | −0.0011 | 0.061 | −0.018 | False | False |
| OFA | RH | subj05 | 883 | 0.0248 | 0.228 | 0.109 | False | False |
| OFA | RH | subj06 | 475 | 0.0134 | 0.124 | 0.108 | False | False |
| OFA | RH | subj07 | 302 | 0.0098 | 0.107 | 0.092 | False | False |
| OFA | RH | subj08 | 231 | −0.0057 | 0.035 | −0.163 | False | False |
| *Place-selective* | | | | | | | | |
| OPA | LH | subj01 | 1863 | 0.0121 | 0.152 | 0.079 | True | True |
| OPA | LH | subj02 | 1494 | 0.0102 | 0.177 | 0.058 | True | True |
| OPA | LH | subj03 | 1797 | 0.0091 | 0.134 | 0.068 | True | True |
| OPA | LH | subj04 | 1707 | 0.0068 | 0.107 | 0.064 | True | True |
| OPA | LH | subj05 | 2203 | 0.0220 | 0.248 | 0.089 | True | True |

*Continued on next page*

Table 1 – *continued from previous page*

| **ROI** | **H** | **Subject** | $N_{\text{vox}}$ | $R^2_{\text{wiped}}$ | $R^2_{\text{raw}}$ | **Ratio** | **Sig.** $t$**-test** | **Sig. SF** |
|---|---|---|---|---|---|---|---|---|
| OPA | LH | subj06 | 1706 | 0.0038 | 0.093 | 0.041 | True | True |
| OPA | LH | subj07 | 1611 | 0.0096 | 0.130 | 0.074 | True | True |
| OPA | LH | subj08 | 2647 | 0.0016 | 0.079 | 0.020 | True | True |
| OPA | RH | subj01 | 2806 | 0.0114 | 0.134 | 0.085 | True | True |
| OPA | RH | subj02 | 2434 | 0.0072 | 0.141 | 0.051 | True | True |
| OPA | RH | subj03 | 1421 | 0.0080 | 0.115 | 0.069 | True | True |
| OPA | RH | subj04 | 1998 | 0.0119 | 0.120 | 0.099 | True | True |
| OPA | RH | subj05 | 2025 | 0.0238 | 0.237 | 0.100 | True | True |
| OPA | RH | subj06 | 1720 | 0.0065 | 0.095 | 0.068 | True | True |
| OPA | RH | subj07 | 1474 | 0.0087 | 0.137 | 0.064 | True | True |
| OPA | RH | subj08 | 1979 | 0.0045 | 0.082 | 0.054 | True | True |
| PPA | LH | subj01 | 1311 | 0.0130 | 0.191 | 0.068 | True | True |
| PPA | LH | subj02 | 1272 | 0.0103 | 0.285 | 0.036 | True | True |
| PPA | LH | subj03 | 1787 | 0.0073 | 0.154 | 0.047 | True | True |
| PPA | LH | subj04 | 1551 | 0.0096 | 0.164 | 0.059 | True | True |
| PPA | LH | subj05 | 1585 | 0.0254 | 0.305 | 0.083 | True | True |
| PPA | LH | subj06 | 1256 | 0.0074 | 0.144 | 0.052 | True | True |
| PPA | LH | subj07 | 1323 | 0.0067 | 0.166 | 0.040 | True | True |
| PPA | LH | subj08 | 1277 | 0.0024 | 0.110 | 0.022 | True | True |
| PPA | RH | subj01 | 891 | 0.0121 | 0.212 | 0.057 | True | True |
| PPA | RH | subj02 | 1490 | 0.0095 | 0.245 | 0.039 | True | True |
| PPA | RH | subj03 | 1976 | 0.0104 | 0.180 | 0.058 | True | True |
| PPA | RH | subj04 | 1314 | 0.0111 | 0.184 | 0.060 | True | True |
| PPA | RH | subj05 | 1586 | 0.0220 | 0.273 | 0.080 | True | True |
| PPA | RH | subj06 | 1512 | 0.0066 | 0.160 | 0.042 | True | True |
| PPA | RH | subj07 | 1466 | 0.0050 | 0.144 | 0.035 | True | True |
| PPA | RH | subj08 | 1326 | 0.0035 | 0.140 | 0.025 | True | True |
| RSC | LH | subj01 | 401 | 0.0154 | 0.268 | 0.058 | True | True |
| RSC | LH | subj02 | 481 | 0.0167 | 0.311 | 0.054 | True | True |
| RSC | LH | subj03 | 515 | 0.0027 | 0.110 | 0.025 | True | True |
| RSC | LH | subj04 | 482 | 0.0094 | 0.158 | 0.059 | True | True |
| RSC | LH | subj05 | 358 | 0.0321 | 0.265 | 0.121 | True | True |
| RSC | LH | subj06 | 323 | 0.0040 | 0.097 | 0.041 | True | True |
| RSC | LH | subj07 | 429 | 0.0016 | 0.081 | 0.020 | True | True |

*Continued on next page*

Table 1 – *continued from previous page*

| **ROI** | **H** | **Subject** | $N_{\text{vox}}$ | $R^2_{\text{wiped}}$ | $R^2_{\text{raw}}$ | **Ratio** | **Sig. $t$-test** | **Sig. SF** |
|---|---|---|---|---|---|---|---|---|
| RSC | LH | subj08 | 339 | 0.0022 | 0.085 | 0.026 | True | True |
| RSC | RH | subj01 | 660 | 0.0103 | 0.206 | 0.050 | True | True |
| RSC | RH | subj02 | 683 | 0.0073 | 0.237 | 0.031 | True | True |
| RSC | RH | subj03 | 890 | 0.0033 | 0.109 | 0.031 | True | True |
| RSC | RH | subj04 | 755 | 0.0128 | 0.176 | 0.073 | True | True |
| RSC | RH | subj05 | 726 | 0.0217 | 0.203 | 0.107 | True | True |
| RSC | RH | subj06 | 687 | 0.0090 | 0.144 | 0.063 | True | True |
| RSC | RH | subj07 | 719 | 0.0147 | 0.178 | 0.083 | True | True |
| RSC | RH | subj08 | 740 | 0.0031 | 0.093 | 0.033 | True | True |
| *Word-selective* | | | | | | | | |
| OWFA | LH | subj01 | 317 | −0.0035 | 0.052 | −0.067 | False | False |
| OWFA | LH | subj02 | 580 | −0.0075 | 0.053 | −0.143 | False | False |
| OWFA | LH | subj03 | 539 | 0.0016 | 0.067 | 0.025 | False | False |
| OWFA | LH | subj04 | 454 | −0.0049 | 0.036 | −0.137 | False | False |
| OWFA | LH | subj05 | 579 | 0.0110 | 0.144 | 0.076 | False | False |
| OWFA | LH | subj06 | 519 | 0.0107 | 0.105 | 0.102 | False | False |
| OWFA | LH | subj07 | 1036 | 0.0012 | 0.068 | 0.018 | False | False |
| OWFA | LH | subj08 | 995 | −0.0059 | 0.036 | −0.161 | False | False |
| OWFA | RH | subj01 | 590 | −0.0040 | 0.035 | −0.115 | False | False |
| OWFA | RH | subj02 | 717 | −0.0052 | 0.045 | −0.116 | False | False |
| OWFA | RH | subj03 | 727 | 0.0009 | 0.050 | 0.017 | False | False |
| OWFA | RH | subj04 | 683 | −0.0041 | 0.040 | −0.104 | False | False |
| OWFA | RH | subj05 | 412 | 0.0179 | 0.190 | 0.094 | False | False |
| OWFA | RH | subj06 | 467 | 0.0048 | 0.072 | 0.067 | False | False |
| OWFA | RH | subj07 | 752 | −0.0064 | 0.022 | −0.291 | False | False |
| OWFA | RH | subj08 | 624 | −0.0053 | 0.037 | −0.144 | False | False |
| VWFA-1 | LH | subj01 | 1381 | 0.0158 | 0.137 | 0.115 | True | True |
| VWFA-1 | LH | subj02 | 499 | 0.0146 | 0.174 | 0.084 | True | True |
| VWFA-1 | LH | subj03 | 1211 | 0.0096 | 0.083 | 0.116 | True | True |
| VWFA-1 | LH | subj04 | 800 | 0.0120 | 0.108 | 0.111 | True | True |
| VWFA-1 | LH | subj05 | 654 | 0.0296 | 0.222 | 0.133 | True | True |
| VWFA-1 | LH | subj06 | 441 | 0.0122 | 0.102 | 0.120 | True | True |
| VWFA-1 | LH | subj07 | 492 | 0.0009 | 0.088 | 0.010 | True | True |
| VWFA-1 | LH | subj08 | 1406 | 0.0065 | 0.059 | 0.110 | True | True |

*Continued on next page*

Table 1 – *continued from previous page*

| **ROI** | **H** | **Subject** | $N_{\text{vox}}$ | $R^2_{\text{wiped}}$ | $R^2_{\text{raw}}$ | **Ratio** | **Sig. $t$-test** | **Sig. SF** |
|---|---|---|---|---|---|---|---|---|
| VWFA-1 | RH | subj01 | 397 | 0.0236 | 0.194 | 0.122 | True | True |
| VWFA-1 | RH | subj02 | 300 | 0.0204 | 0.214 | 0.095 | True | True |
| VWFA-1 | RH | subj03 | 502 | 0.0139 | 0.106 | 0.131 | True | True |
| VWFA-1 | RH | subj04 | 644 | 0.0146 | 0.136 | 0.107 | True | True |
| VWFA-1 | RH | subj05 | 313 | 0.0194 | 0.173 | 0.112 | True | True |
| VWFA-1 | RH | subj06 | 456 | 0.0160 | 0.122 | 0.131 | True | True |
| VWFA-1 | RH | subj07 | 416 | 0.0281 | 0.233 | 0.120 | True | True |
| VWFA-1 | RH | subj08 | 639 | 0.0089 | 0.072 | 0.123 | True | True |
| VWFA-2 | LH | subj01 | 461 | 0.0158 | 0.109 | 0.144 | True | True |
| VWFA-2 | LH | subj02 | 189 | 0.0220 | 0.139 | 0.159 | True | True |
| VWFA-2 | LH | subj03 | 36 | 0.0329 | 0.109 | 0.302 | True | True |
| VWFA-2 | LH | subj04 | 46 | 0.0367 | 0.138 | 0.266 | True | True |
| VWFA-2 | LH | subj05 | 329 | 0.0307 | 0.190 | 0.162 | True | True |
| VWFA-2 | LH | subj06 | 678 | 0.0123 | 0.062 | 0.199 | True | True |
| VWFA-2 | LH | subj07 | 632 | 0.0199 | 0.132 | 0.151 | True | True |
| VWFA-2 | LH | subj08 | 126 | −0.0003 | 0.033 | −0.010 | True | True |
| VWFA-2 | RH | subj01 | 431 | 0.0165 | 0.123 | 0.135 | True | False |
| VWFA-2 | RH | subj02 | 155 | 0.0150 | 0.161 | 0.093 | True | False |
| VWFA-2 | RH | subj05 | 338 | 0.0138 | 0.148 | 0.093 | True | False |
| VWFA-2 | RH | subj06 | 186 | 0.0224 | 0.127 | 0.177 | True | False |
| VWFA-2 | RH | subj07 | 1107 | 0.0180 | 0.147 | 0.122 | True | False |
| mTL-words | RH | subj07 | 63 | −0.0004 | 0.052 | −0.008 | False | False |
| mfs-words | LH | subj01 | 490 | 0.0117 | 0.101 | 0.116 | False | True |
| mfs-words | LH | subj02 | 366 | 0.0051 | 0.141 | 0.036 | False | True |
| mfs-words | LH | subj03 | 79 | 0.0074 | 0.110 | 0.067 | False | True |
| mfs-words | LH | subj04 | 146 | 0.0104 | 0.105 | 0.098 | False | True |
| mfs-words | LH | subj05 | 257 | 0.0450 | 0.318 | 0.141 | False | True |
| mfs-words | LH | subj06 | 432 | 0.0148 | 0.106 | 0.140 | False | True |
| mfs-words | LH | subj07 | 182 | 0.0039 | 0.094 | 0.041 | False | True |
| mfs-words | LH | subj08 | 146 | −0.0027 | 0.040 | −0.067 | False | True |
| mfs-words | RH | subj02 | 400 | 0.0042 | 0.109 | 0.039 | False | False |
| mfs-words | RH | subj04 | 106 | 0.0107 | 0.123 | 0.087 | False | False |
| mfs-words | RH | subj05 | 340 | 0.0269 | 0.223 | 0.120 | False | False |
| mfs-words | RH | subj06 | 431 | 0.0139 | 0.110 | 0.127 | False | False |

*Continued on next page*

Table 1 – *continued from previous page*

| **ROI** | **H** | **Subject** | $N_{\text{vox}}$ | $R^2_{\text{wiped}}$ | $R^2_{\text{raw}}$ | **Ratio** | **Sig.** $t$**-test** | **Sig. SF** |
|---|---|---|---|---|---|---|---|---|
| mfs-words | RH | subj07 | 298 | 0.0070 | 0.093 | 0.075 | False | False |
| mfs-words | RH | subj08 | 222 | −0.0003 | 0.075 | −0.004 | False | False |
| *Early visual (retinotopic)* | | | | | | | | |
| V1d | LH | subj01 | 828 | −0.0168 | 0.046 | −0.370 | True | False |
| V1d | LH | subj02 | 845 | −0.0165 | 0.081 | −0.205 | True | False |
| V1d | LH | subj03 | 523 | −0.0145 | 0.063 | −0.229 | True | False |
| V1d | LH | subj04 | 557 | −0.0123 | 0.059 | −0.207 | True | False |
| V1d | LH | subj05 | 1117 | −0.0101 | 0.101 | −0.100 | True | False |
| V1d | LH | subj06 | 944 | −0.0118 | 0.053 | −0.221 | True | False |
| V1d | LH | subj07 | 917 | −0.0140 | 0.047 | −0.298 | True | False |
| V1d | LH | subj08 | 569 | −0.0146 | 0.067 | −0.219 | True | False |
| V1d | RH | subj01 | 991 | −0.0156 | 0.040 | −0.390 | True | False |
| V1d | RH | subj02 | 677 | −0.0167 | 0.078 | −0.213 | True | False |
| V1d | RH | subj03 | 624 | −0.0127 | 0.062 | −0.206 | True | False |
| V1d | RH | subj04 | 553 | −0.0124 | 0.068 | −0.181 | True | False |
| V1d | RH | subj05 | 706 | −0.0131 | 0.103 | −0.127 | True | False |
| V1d | RH | subj06 | 712 | −0.0140 | 0.067 | −0.208 | True | False |
| V1d | RH | subj07 | 709 | −0.0145 | 0.047 | −0.309 | True | False |
| V1d | RH | subj08 | 815 | −0.0135 | 0.047 | −0.289 | True | False |
| V1v | LH | subj01 | 710 | −0.0167 | 0.043 | −0.384 | True | False |
| V1v | LH | subj02 | 599 | −0.0157 | 0.046 | −0.343 | True | False |
| V1v | LH | subj03 | 853 | −0.0154 | 0.063 | −0.246 | True | False |
| V1v | LH | subj04 | 585 | −0.0132 | 0.068 | −0.195 | True | False |
| V1v | LH | subj05 | 538 | −0.0130 | 0.085 | −0.153 | True | False |
| V1v | LH | subj06 | 299 | −0.0141 | 0.073 | −0.194 | True | False |
| V1v | LH | subj07 | 491 | −0.0129 | 0.048 | −0.270 | True | False |
| V1v | LH | subj08 | 444 | −0.0156 | 0.057 | −0.273 | True | False |
| V1v | RH | subj01 | 444 | −0.0155 | 0.038 | −0.407 | True | False |
| V1v | RH | subj02 | 616 | −0.0184 | 0.065 | −0.283 | True | False |
| V1v | RH | subj03 | 676 | −0.0127 | 0.053 | −0.240 | True | False |
| V1v | RH | subj04 | 633 | −0.0127 | 0.077 | −0.165 | True | False |
| V1v | RH | subj05 | 589 | −0.0139 | 0.127 | −0.109 | True | False |
| V1v | RH | subj06 | 572 | −0.0133 | 0.061 | −0.218 | True | False |
| V1v | RH | subj07 | 444 | −0.0135 | 0.051 | −0.266 | True | False |

*Continued on next page*

Table 1 – *continued from previous page*

| **ROI** | **H** | **Subject** | $N_{\text{vox}}$ | $R^2_{\text{wiped}}$ | $R^2_{\text{raw}}$ | **Ratio** | **Sig.** $t$**-test** | **Sig. SF** |
|---|---|---|---|---|---|---|---|---|
| V1v | RH | subj08 | 590 | −0.0123 | 0.033 | −0.371 | True | False |
| V2d | LH | subj01 | 692 | −0.0137 | 0.037 | −0.371 | True | False |
| V2d | LH | subj02 | 620 | −0.0109 | 0.038 | −0.284 | True | False |
| V2d | LH | subj03 | 575 | −0.0095 | 0.046 | −0.206 | True | False |
| V2d | LH | subj04 | 498 | −0.0144 | 0.051 | −0.284 | True | False |
| V2d | LH | subj05 | 663 | −0.0067 | 0.082 | −0.082 | True | False |
| V2d | LH | subj06 | 557 | −0.0081 | 0.064 | −0.125 | True | False |
| V2d | LH | subj07 | 616 | −0.0096 | 0.037 | −0.260 | True | False |
| V2d | LH | subj08 | 503 | −0.0114 | 0.032 | −0.355 | True | False |
| V2d | RH | subj01 | 725 | −0.0137 | 0.051 | −0.269 | True | False |
| V2d | RH | subj02 | 623 | −0.0114 | 0.042 | −0.271 | True | False |
| V2d | RH | subj03 | 782 | −0.0094 | 0.047 | −0.198 | True | False |
| V2d | RH | subj04 | 595 | −0.0122 | 0.044 | −0.276 | True | False |
| V2d | RH | subj05 | 796 | −0.0074 | 0.068 | −0.109 | True | False |
| V2d | RH | subj06 | 559 | −0.0098 | 0.076 | −0.129 | True | False |
| V2d | RH | subj07 | 523 | −0.0110 | 0.036 | −0.307 | True | False |
| V2d | RH | subj08 | 719 | −0.0103 | 0.040 | −0.256 | True | False |
| V2v | LH | subj01 | 632 | −0.0145 | 0.042 | −0.344 | True | False |
| V2v | LH | subj02 | 653 | −0.0146 | 0.047 | −0.314 | True | False |
| V2v | LH | subj03 | 930 | −0.0115 | 0.069 | −0.165 | True | False |
| V2v | LH | subj04 | 734 | −0.0122 | 0.069 | −0.178 | True | False |
| V2v | LH | subj05 | 531 | −0.0086 | 0.121 | −0.072 | True | False |
| V2v | LH | subj06 | 482 | −0.0114 | 0.064 | −0.179 | True | False |
| V2v | LH | subj07 | 685 | −0.0119 | 0.058 | −0.204 | True | False |
| V2v | LH | subj08 | 646 | −0.0124 | 0.064 | −0.194 | True | False |
| V2v | RH | subj01 | 887 | −0.0140 | 0.038 | −0.369 | True | False |
| V2v | RH | subj02 | 883 | −0.0125 | 0.030 | −0.424 | True | False |
| V2v | RH | subj03 | 704 | −0.0085 | 0.050 | −0.170 | True | False |
| V2v | RH | subj04 | 647 | −0.0130 | 0.070 | −0.185 | True | False |
| V2v | RH | subj05 | 629 | −0.0069 | 0.114 | −0.060 | True | False |
| V2v | RH | subj06 | 610 | −0.0095 | 0.063 | −0.150 | True | False |
| V2v | RH | subj07 | 575 | −0.0112 | 0.065 | −0.174 | True | False |
| V2v | RH | subj08 | 744 | −0.0116 | 0.051 | −0.227 | True | False |
| V3d | LH | subj01 | 669 | −0.0106 | 0.053 | −0.199 | True | False |

*Continued on next page*

Table 1 – *continued from previous page*

| **ROI** | **H** | **Subject** | $N_{\text{vox}}$ | $R^2_{\text{wiped}}$ | $R^2_{\text{raw}}$ | **Ratio** | **Sig.** $t$**-test** | **Sig. SF** |
|---|---|---|---|---|---|---|---|---|
| V3d | LH | subj02 | 574 | −0.0118 | 0.058 | −0.204 | True | False |
| V3d | LH | subj03 | 653 | −0.0057 | 0.043 | −0.132 | True | False |
| V3d | LH | subj04 | 468 | −0.0102 | 0.041 | −0.252 | True | False |
| V3d | LH | subj05 | 725 | −0.0009 | 0.096 | −0.009 | True | False |
| V3d | LH | subj06 | 622 | −0.0057 | 0.043 | −0.133 | True | False |
| V3d | LH | subj07 | 651 | −0.0084 | 0.034 | −0.249 | True | False |
| V3d | LH | subj08 | 489 | −0.0097 | 0.036 | −0.270 | True | False |
| V3d | RH | subj01 | 535 | −0.0104 | 0.058 | −0.179 | True | False |
| V3d | RH | subj02 | 756 | −0.0110 | 0.048 | −0.229 | True | False |
| V3d | RH | subj03 | 676 | −0.0070 | 0.049 | −0.143 | True | False |
| V3d | RH | subj04 | 578 | −0.0095 | 0.040 | −0.238 | True | False |
| V3d | RH | subj05 | 714 | −0.0029 | 0.085 | −0.034 | True | False |
| V3d | RH | subj06 | 527 | −0.0060 | 0.053 | −0.113 | True | False |
| V3d | RH | subj07 | 550 | −0.0089 | 0.039 | −0.228 | True | False |
| V3d | RH | subj08 | 766 | −0.0085 | 0.048 | −0.177 | True | False |
| V3v | LH | subj01 | 567 | −0.0113 | 0.048 | −0.233 | True | False |
| V3v | LH | subj02 | 553 | −0.0123 | 0.039 | −0.315 | True | False |
| V3v | LH | subj03 | 656 | −0.0047 | 0.066 | −0.071 | True | False |
| V3v | LH | subj04 | 537 | −0.0082 | 0.047 | −0.173 | True | False |
| V3v | LH | subj05 | 457 | −0.0008 | 0.117 | −0.007 | True | False |
| V3v | LH | subj06 | 422 | −0.0098 | 0.051 | −0.193 | True | False |
| V3v | LH | subj07 | 489 | −0.0097 | 0.055 | −0.176 | True | False |
| V3v | LH | subj08 | 509 | −0.0091 | 0.062 | −0.145 | True | False |
| V3v | RH | subj01 | 682 | −0.0112 | 0.046 | −0.246 | True | False |
| V3v | RH | subj02 | 732 | −0.0136 | 0.059 | −0.229 | True | False |
| V3v | RH | subj03 | 433 | −0.0039 | 0.054 | −0.073 | True | False |
| V3v | RH | subj04 | 563 | −0.0086 | 0.050 | −0.171 | True | False |
| V3v | RH | subj05 | 487 | −0.0017 | 0.081 | −0.021 | True | False |
| V3v | RH | subj06 | 532 | −0.0070 | 0.054 | −0.129 | True | False |
| V3v | RH | subj07 | 570 | −0.0074 | 0.068 | −0.108 | True | False |
| V3v | RH | subj08 | 527 | −0.0090 | 0.044 | −0.202 | True | False |
| hV4 | LH | subj01 | 531 | −0.0074 | 0.055 | −0.135 | False | False |
| hV4 | LH | subj02 | 606 | −0.0099 | 0.085 | −0.117 | False | False |
| hV4 | LH | subj03 | 457 | 0.0005 | 0.081 | 0.006 | False | False |

*Continued on next page*

Table 1 – *continued from previous page*

| **ROI** | **H** | **Subject** | $N_{\text{vox}}$ | $R^2_{\text{wiped}}$ | $R^2_{\text{raw}}$ | **Ratio** | **Sig. $t$-test** | **Sig. SF** |
|---|---|---|---|---|---|---|---|---|
| hV4 | LH | subj04 | 547 | −0.0051 | 0.060 | −0.085 | False | False |
| hV4 | LH | subj05 | 584 | 0.0073 | 0.145 | 0.050 | False | False |
| hV4 | LH | subj06 | 439 | −0.0009 | 0.071 | −0.012 | False | False |
| hV4 | LH | subj07 | 540 | −0.0054 | 0.059 | −0.091 | False | False |
| hV4 | LH | subj08 | 538 | −0.0077 | 0.076 | −0.102 | False | False |
| hV4 | RH | subj01 | 765 | −0.0069 | 0.080 | −0.086 | False | False |
| hV4 | RH | subj02 | 656 | −0.0100 | 0.083 | −0.121 | False | False |
| hV4 | RH | subj03 | 430 | −0.0001 | 0.065 | −0.002 | False | False |
| hV4 | RH | subj04 | 643 | −0.0038 | 0.062 | −0.061 | False | False |
| hV4 | RH | subj05 | 336 | 0.0046 | 0.108 | 0.042 | False | False |
| hV4 | RH | subj06 | 376 | −0.0020 | 0.057 | −0.035 | False | False |
| hV4 | RH | subj07 | 381 | −0.0059 | 0.044 | −0.136 | False | False |
| hV4 | RH | subj08 | 538 | −0.0085 | 0.062 | −0.136 | False | False |

H = hemisphere; Subject = participant ID; $N_{\text{vox}}$ = number of voxels in ROI for that subject.
$R^2_{\text{wiped}}$ = visually-independent semantic encoding; $R^2_{\text{raw}}$ = raw encoding; Ratio = $R^2_{\text{wiped}}/R^2_{\text{raw}}$.
Sig. $t$-test: group-level significance by one-sample $t$-test (BH-FDR corrected).
Sig. SF: group-level significance by sign-flip permutation test (BH-FDR corrected).

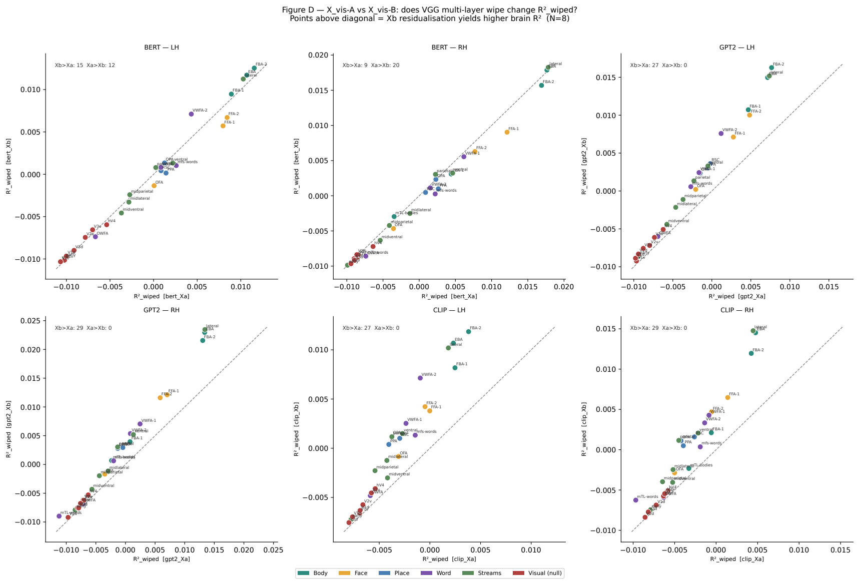


Figure S1: **Supplementary Figure S1. $\mathbf{X}_{\mathrm{vis}}$-A vs. $\mathbf{X}_{\mathrm{vis}}$-B residualisation: effect on $R^2_{\mathrm{wiped}}$ across cortical ROIs.** Each panel plots $R^2_{\mathrm{wiped}}$ under $\mathrm{X}_{\mathrm{vis}}$-A residualisation (broad ensemble: Gabor + VGG19 + ResNet50 + CLIP-visual + EfficientNet-B0; $x$-axis) against $\mathrm{X}_{\mathrm{vis}}$-B (VGG19 hierarchical multi-layer GAP; $y$-axis) for three language models (BERT, GPT-2, CLIP-text) × two hemispheres ($N = 8$). Points are individual ROIs coloured by functional group; the dashed line is the identity. Pearson correlations between conditions are uniformly high ($r = 0.977$–$0.998$, all $p < 10^{-17}$), confirming that visual feature set choice does not alter ROI rank ordering. GPT-2 and CLIP show a systematic above-diagonal shift ($\mathrm{X}_{\mathrm{vis}}\text{-B} > \mathrm{X}_{\mathrm{vis}}\text{-A}$ in 27–29 of 27–28 ROIs per hemisphere), whereas BERT shows no consistent directional preference (LH: 15 vs. 12; RH: 9 vs. 20). Deviations from the diagonal are small in all cases (mean $|\mathrm{X}_{\mathrm{vis}}\text{-B} - \mathrm{X}_{\mathrm{vis}}\text{-A}| < 0.001$), supporting the equivalence of both residualisation strategies for the main analysis.

Table 2: **Group-level visually-independent semantic encoding ($R^2_{\text{wiped}}$) across visual ROIs for all models and variants.** Values are mean ± SEM across subjects. Ratio = $R^2_{\text{wiped}}/R^2_{\text{raw}}$. $g$ = Hedges' $g$ (effect size). $q_{\text{FDR}}$: BH-corrected $p$-value from one-sample $t$-test. $q_{\text{SF}}$: BH-corrected sign-flip permutation $p$-value. Rows are grouped by ROI category, sorted by ROI, hemisphere, then model.

| **ROI** | **H** | **Model** | **N** | $R^2_{\text{wiped}}$ **(mean ± SEM)** | $R^2_{\text{raw}}$ | **Ratio** | $g$ | $q_{\text{FDR}}$ | $q_{\text{SF}}$ |
|---|---|---|---|---|---|---|---|---|---|
| *Stream-level ROIs* | | | | | | | | | |
| Early stream | LH | bert_Xa | 8 | $-0.0102 \pm 0.0009$ | 0.048 | −0.234 | -3.74 | <0.001 | 1.000 |
| Early stream | LH | bert_Xb | 8 | $-0.0099 \pm 0.0008$ | 0.048 | −0.227 | -3.78 | <0.001 | 1.000 |
| Early stream | LH | gpt2_Xa | 8 | $-0.0094 \pm 0.0008$ | 0.048 | −0.214 | -3.94 | <0.001 | 1.000 |
| Early stream | LH | gpt2_Xb | 8 | $-0.0086 \pm 0.0008$ | 0.048 | −0.197 | -3.40 | <0.001 | 1.000 |
| Early stream | LH | clip_Xa | 8 | $-0.0078 \pm 0.0006$ | 0.048 | −0.174 | -3.86 | <0.001 | 1.000 |
| Early stream | LH | clip_Xb | 8 | $-0.0072 \pm 0.0006$ | 0.048 | −0.163 | -3.99 | <0.001 | 1.000 |
| Early stream | RH | bert_Xa | 8 | $-0.0099 \pm 0.0010$ | 0.047 | −0.234 | -3.27 | <0.001 | 1.000 |
| Early stream | RH | bert_Xb | 8 | $-0.0099 \pm 0.0010$ | 0.047 | −0.234 | -3.11 | <0.001 | 1.000 |
| Early stream | RH | gpt2_Xa | 8 | $-0.0085 \pm 0.0007$ | 0.047 | −0.199 | -4.07 | <0.001 | 1.000 |
| Early stream | RH | gpt2_Xb | 8 | $-0.0079 \pm 0.0007$ | 0.047 | −0.186 | -3.49 | <0.001 | 1.000 |
| Early stream | RH | clip_Xa | 8 | $-0.0079 \pm 0.0006$ | 0.047 | −0.181 | -4.19 | <0.001 | 1.000 |
| Early stream | RH | clip_Xb | 8 | $-0.0075 \pm 0.0005$ | 0.047 | −0.173 | -4.75 | <0.001 | 1.000 |
| Lateral stream | LH | bert_Xa | 8 | $0.0103 \pm 0.0027$ | 0.175 | 0.052 | 1.21 | 0.015 | 0.027* |
| Lateral stream | LH | bert_Xb | 8 | $0.0112 \pm 0.0031$ | 0.175 | 0.056 | 1.14 | 0.020 | 0.043* |
| Lateral stream | LH | gpt2_Xa | 8 | $0.0074 \pm 0.0024$ | 0.175 | 0.035 | 0.97 | 0.038 | 0.107 |
| Lateral stream | LH | gpt2_Xb | 8 | $0.0152 \pm 0.0034$ | 0.175 | 0.079 | 1.41 | 0.008 | 0.027* |

*Continued on next page*

Table 2 – *continued from previous page*

| ROI | H | Model | N | $R^2_{\text{wiped}}$ (mean ± SEM) | $R^2_{\text{raw}}$ | Ratio | $g$ | $q_{\text{FDR}}$ | $q_{\text{SF}}$ |
|---|---|---|---|---|---|---|---|---|---|
| Lateral stream | LH | clip_Xa | 8 | $0.0018 \pm 0.0010$ | 0.175 | 0.007 | 0.57 | 0.155 | 0.501 |
| Lateral stream | LH | clip_Xb | 8 | $0.0102 \pm 0.0024$ | 0.175 | 0.053 | 1.35 | 0.010 | 0.031* |
| Lateral stream | RH | bert_Xa | 8 | $0.0178 \pm 0.0030$ | 0.219 | 0.077 | 1.85 | 0.002 | 0.027* |
| Lateral stream | RH | bert_Xb | 8 | $0.0183 \pm 0.0037$ | 0.219 | 0.077 | 1.57 | 0.005 | 0.043* |
| Lateral stream | RH | gpt2_Xa | 8 | $0.0134 \pm 0.0024$ | 0.219 | 0.057 | 1.77 | 0.003 | 0.054 |
| Lateral stream | RH | gpt2_Xb | 8 | $0.0234 \pm 0.0037$ | 0.219 | 0.101 | 1.97 | 0.002 | 0.027* |
| Lateral stream | RH | clip_Xa | 8 | $0.0045 \pm 0.0012$ | 0.219 | 0.018 | 1.17 | 0.015 | 0.215 |
| Lateral stream | RH | clip_Xb | 8 | $0.0148 \pm 0.0029$ | 0.219 | 0.063 | 1.59 | 0.006 | 0.031* |
| Mid-lateral | LH | bert_Xa | 8 | $-0.0028 \pm 0.0016$ | 0.094 | $-0.040$ | -0.54 | 0.209 | 1.000 |
| Mid-lateral | LH | bert_Xb | 8 | $-0.0033 \pm 0.0015$ | 0.094 | $-0.044$ | -0.67 | 0.118 | 1.000 |
| Mid-lateral | LH | gpt2_Xa | 8 | $-0.0046 \pm 0.0015$ | 0.094 | $-0.059$ | -0.94 | 0.041 | 1.000 |
| Mid-lateral | LH | gpt2_Xb | 8 | $-0.0022 \pm 0.0019$ | 0.094 | $-0.032$ | -0.37 | 0.345 | 1.000 |
| Mid-lateral | LH | clip_Xa | 8 | $-0.0042 \pm 0.0008$ | 0.094 | $-0.049$ | -1.75 | 0.002 | 1.000 |
| Mid-lateral | LH | clip_Xb | 8 | $-0.0012 \pm 0.0010$ | 0.094 | $-0.016$ | -0.39 | 0.349 | 1.000 |
| Mid-lateral | RH | bert_Xa | 8 | $-0.0013 \pm 0.0021$ | 0.103 | $-0.040$ | -0.19 | 0.639 | 1.000 |
| Mid-lateral | RH | bert_Xb | 8 | $-0.0025 \pm 0.0020$ | 0.103 | $-0.053$ | -0.39 | 0.357 | 1.000 |
| Mid-lateral | RH | gpt2_Xa | 8 | $-0.0029 \pm 0.0018$ | 0.103 | $-0.052$ | -0.52 | 0.228 | 1.000 |
| Mid-lateral | RH | gpt2_Xb | 8 | $-0.0011 \pm 0.0023$ | 0.103 | $-0.037$ | -0.16 | 0.678 | 1.000 |
| Mid-lateral | RH | clip_Xa | 8 | $-0.0052 \pm 0.0012$ | 0.103 | $-0.066$ | -1.42 | 0.006 | 1.000 |
| Mid-lateral | RH | clip_Xb | 8 | $-0.0025 \pm 0.0016$ | 0.103 | $-0.043$ | -0.49 | 0.244 | 1.000 |
| Mid-parietal | LH | bert_Xa | 8 | $-0.0027 \pm 0.0020$ | 0.098 | $-0.050$ | -0.42 | 0.317 | 1.000 |
| Mid-parietal | LH | bert_Xb | 8 | $-0.0024 \pm 0.0023$ | 0.098 | $-0.047$ | -0.34 | 0.430 | 1.000 |
| Mid-parietal | LH | gpt2_Xa | 8 | $-0.0037 \pm 0.0026$ | 0.098 | $-0.065$ | -0.45 | 0.292 | 1.000 |
| Mid-parietal | LH | gpt2_Xb | 8 | $-0.0011 \pm 0.0032$ | 0.098 | $-0.040$ | -0.11 | 0.780 | 1.000 |
| Mid-parietal | LH | clip_Xa | 8 | $-0.0054 \pm 0.0010$ | 0.098 | $-0.073$ | -1.78 | 0.002 | 1.000 |
| Mid-parietal | LH | clip_Xb | 8 | $-0.0023 \pm 0.0018$ | 0.098 | $-0.042$ | -0.40 | 0.345 | 1.000 |

*Continued on next page*

Table 2 – *continued from previous page*

| **ROI** | **H** | **Model** | **N** | $R^2_{\text{wiped}}$ **(mean ± SEM)** | $R^2_{\text{raw}}$ | **Ratio** | $g$ | $q_{\text{FDR}}$ | $q_{\text{SF}}$ |
|---|---|---|---|---|---|---|---|---|---|
| Mid-parietal | RH | bert_Xa | 8 | $-0.0041 \pm 0.0014$ | 0.085 | −0.065 | -0.95 | 0.036 | 1.000 |
| Mid-parietal | RH | bert_Xb | 8 | $-0.0042 \pm 0.0014$ | 0.085 | −0.065 | -0.94 | 0.039 | 1.000 |
| Mid-parietal | RH | gpt2_Xa | 8 | $-0.0044 \pm 0.0016$ | 0.085 | −0.072 | -0.87 | 0.053 | 1.000 |
| Mid-parietal | RH | gpt2_Xb | 8 | $-0.0020 \pm 0.0021$ | 0.085 | −0.042 | -0.29 | 0.427 | 1.000 |
| Mid-parietal | RH | clip_Xa | 8 | $-0.0065 \pm 0.0010$ | 0.085 | −0.095 | -2.04 | 0.001 | 1.000 |
| Mid-parietal | RH | clip_Xb | 8 | $-0.0040 \pm 0.0010$ | 0.085 | −0.063 | -1.21 | 0.015 | 1.000 |
| Mid-ventral | LH | bert_Xa | 8 | $-0.0037 \pm 0.0012$ | 0.088 | −0.055 | -0.96 | 0.036 | 1.000 |
| Mid-ventral | LH | bert_Xb | 8 | $-0.0046 \pm 0.0010$ | 0.088 | −0.064 | -1.46 | 0.007 | 1.000 |
| Mid-ventral | LH | gpt2_Xa | 8 | $-0.0058 \pm 0.0012$ | 0.088 | −0.079 | -1.49 | 0.006 | 1.000 |
| Mid-ventral | LH | gpt2_Xb | 8 | $-0.0044 \pm 0.0015$ | 0.088 | −0.063 | -0.95 | 0.039 | 1.000 |
| Mid-ventral | LH | clip_Xa | 8 | $-0.0042 \pm 0.0008$ | 0.088 | −0.057 | -1.75 | 0.002 | 1.000 |
| Mid-ventral | LH | clip_Xb | 8 | $-0.0030 \pm 0.0010$ | 0.088 | −0.043 | -0.93 | 0.039 | 1.000 |
| Mid-ventral | RH | bert_Xa | 8 | $-0.0054 \pm 0.0017$ | 0.083 | −0.094 | -0.99 | 0.033 | 1.000 |
| Mid-ventral | RH | bert_Xb | 8 | $-0.0063 \pm 0.0016$ | 0.083 | −0.107 | -1.22 | 0.017 | 1.000 |
| Mid-ventral | RH | gpt2_Xa | 8 | $-0.0057 \pm 0.0013$ | 0.083 | −0.093 | -1.39 | 0.008 | 1.000 |
| Mid-ventral | RH | gpt2_Xb | 8 | $-0.0043 \pm 0.0016$ | 0.083 | −0.078 | -0.83 | 0.057 | 1.000 |
| Mid-ventral | RH | clip_Xa | 8 | $-0.0052 \pm 0.0009$ | 0.083 | −0.083 | -1.80 | 0.002 | 1.000 |
| Mid-ventral | RH | clip_Xb | 8 | $-0.0041 \pm 0.0013$ | 0.083 | −0.070 | -0.95 | 0.037 | 1.000 |
| Parietal stream | LH | bert_Xa | 8 | $0.0003 \pm 0.0011$ | 0.105 | −0.005 | 0.07 | 0.845 | 0.828 |
| Parietal stream | LH | bert_Xb | 8 | $0.0008 \pm 0.0010$ | 0.105 | 0.001 | 0.24 | 0.568 | 0.704 |
| Parietal stream | LH | gpt2_Xa | 8 | $-0.0023 \pm 0.0009$ | 0.105 | −0.034 | -0.76 | 0.085 | 1.000 |
| Parietal stream | LH | gpt2_Xb | 8 | $0.0013 \pm 0.0012$ | 0.105 | 0.002 | 0.35 | 0.365 | 0.358 |
| Parietal stream | LH | clip_Xa | 8 | $-0.0038 \pm 0.0013$ | 0.105 | −0.038 | -0.93 | 0.038 | 1.000 |
| Parietal stream | LH | clip_Xb | 8 | $0.0012 \pm 0.0011$ | 0.105 | 0.012 | 0.33 | 0.414 | 0.493 |
| Parietal stream | RH | bert_Xa | 8 | $0.0022 \pm 0.0020$ | 0.103 | 0.014 | 0.35 | 0.385 | 0.513 |
| Parietal stream | RH | bert_Xb | 8 | $0.0030 \pm 0.0022$ | 0.103 | 0.021 | 0.44 | 0.307 | 0.602 |

*Continued on next page*

Table 2 – *continued from previous page*

| **ROI** | **H** | **Model** | **N** | $R^2_{\text{wiped}}$ **(mean ± SEM)** | $R^2_{\text{raw}}$ | **Ratio** | $g$ | $q_{\text{FDR}}$ | $q_{\text{SF}}$ |
|---|---|---|---|---|---|---|---|---|---|
| Parietal stream | RH | gpt2_Xa | 8 | $-0.0013 \pm 0.0015$ | 0.103 | $-0.030$ | -0.28 | 0.494 | 1.000 |
| Parietal stream | RH | gpt2_Xb | 8 | $0.0030 \pm 0.0022$ | 0.103 | 0.015 | 0.44 | 0.275 | 0.303 |
| Parietal stream | RH | clip_Xa | 8 | $-0.0045 \pm 0.0013$ | 0.103 | $-0.052$ | -1.10 | 0.019 | 1.000 |
| Parietal stream | RH | clip_Xb | 8 | $0.0012 \pm 0.0015$ | 0.103 | 0.006 | 0.25 | 0.514 | 0.560 |
| Ventral stream | LH | bert_Xa | 8 | $0.0022 \pm 0.0020$ | 0.149 | 0.008 | 0.35 | 0.385 | 0.543 |
| Ventral stream | LH | bert_Xb | 8 | $0.0013 \pm 0.0019$ | 0.149 | 0.001 | 0.21 | 0.607 | 0.908 |
| Ventral stream | LH | gpt2_Xa | 8 | $-0.0005 \pm 0.0016$ | 0.149 | $-0.012$ | -0.10 | 0.808 | 1.000 |
| Ventral stream | LH | gpt2_Xb | 8 | $0.0033 \pm 0.0022$ | 0.149 | 0.013 | 0.48 | 0.245 | 0.215 |
| Ventral stream | LH | clip_Xa | 8 | $-0.0027 \pm 0.0011$ | 0.149 | $-0.023$ | -0.75 | 0.074 | 1.000 |
| Ventral stream | LH | clip_Xb | 8 | $0.0015 \pm 0.0016$ | 0.149 | 0.005 | 0.29 | 0.474 | 0.557 |
| Ventral stream | RH | bert_Xa | 8 | $0.0046 \pm 0.0011$ | 0.161 | 0.028 | 1.33 | 0.011 | 0.027* |
| Ventral stream | RH | bert_Xb | 8 | $0.0032 \pm 0.0012$ | 0.161 | 0.019 | 0.85 | 0.055 | 0.061 |
| Ventral stream | RH | gpt2_Xa | 8 | $0.0013 \pm 0.0010$ | 0.161 | 0.006 | 0.44 | 0.302 | 0.537 |
| Ventral stream | RH | gpt2_Xb | 8 | $0.0052 \pm 0.0013$ | 0.161 | 0.030 | 1.30 | 0.011 | 0.027* |
| Ventral stream | RH | clip_Xa | 8 | $-0.0022 \pm 0.0009$ | 0.161 | $-0.013$ | -0.72 | 0.080 | 1.000 |
| Ventral stream | RH | clip_Xb | 8 | $0.0021 \pm 0.0009$ | 0.161 | 0.013 | 0.70 | 0.108 | 0.150 |
| *Body-selective* | | | | | | | | | |
| EBA | LH | bert_Xa | 8 | $0.0107 \pm 0.0026$ | 0.176 | 0.056 | 1.30 | 0.011 | 0.027* |
| EBA | LH | bert_Xb | 8 | $0.0117 \pm 0.0031$ | 0.176 | 0.060 | 1.18 | 0.019 | 0.043* |
| EBA | LH | gpt2_Xa | 8 | $0.0071 \pm 0.0025$ | 0.176 | 0.033 | 0.90 | 0.049 | 0.107 |
| EBA | LH | gpt2_Xb | 8 | $0.0150 \pm 0.0036$ | 0.176 | 0.077 | 1.29 | 0.011 | 0.027* |
| EBA | LH | clip_Xa | 8 | $0.0023 \pm 0.0010$ | 0.176 | 0.011 | 0.70 | 0.086 | 0.344 |
| EBA | LH | clip_Xb | 8 | $0.0107 \pm 0.0025$ | 0.176 | 0.056 | 1.36 | 0.010 | 0.031* |
| EBA | RH | bert_Xa | 8 | $0.0176 \pm 0.0026$ | 0.222 | 0.077 | 2.12 | 0.001 | 0.027* |
| EBA | RH | bert_Xb | 8 | $0.0179 \pm 0.0032$ | 0.222 | 0.076 | 1.75 | 0.003 | 0.043* |

*Continued on next page*

Table 2 – *continued from previous page*

| **ROI** | **H** | **Model** | **N** | $R^2_{\text{wiped}}$ **(mean ± SEM)** | $R^2_{\text{raw}}$ | **Ratio** | $g$ | $q_{\text{FDR}}$ | $q_{\text{SF}}$ |
|---|---|---|---|---|---|---|---|---|---|
| EBA | RH | gpt2_Xa | 8 | $0.0134 \pm 0.0024$ | 0.222 | 0.056 | 1.75 | 0.003 | 0.054 |
| EBA | RH | gpt2_Xb | 8 | $0.0230 \pm 0.0036$ | 0.222 | 0.098 | 2.02 | 0.002 | 0.027* |
| EBA | RH | clip_Xa | 8 | $0.0048 \pm 0.0011$ | 0.222 | 0.020 | 1.39 | 0.007 | 0.215 |
| EBA | RH | clip_Xb | 8 | $0.0145 \pm 0.0025$ | 0.222 | 0.062 | 1.84 | 0.003 | 0.031* |
| FBA-1 | LH | bert_Xa | 5 | $0.0089 \pm 0.0028$ | 0.151 | 0.060 | 1.15 | 0.056 | 0.156 |
| FBA-1 | LH | bert_Xb | 5 | $0.0095 \pm 0.0032$ | 0.151 | 0.066 | 1.07 | 0.071 | 0.286 |
| FBA-1 | LH | gpt2_Xa | 5 | $0.0047 \pm 0.0028$ | 0.151 | 0.028 | 0.59 | 0.272 | 0.537 |
| FBA-1 | LH | gpt2_Xb | 5 | $0.0107 \pm 0.0041$ | 0.151 | 0.069 | 0.93 | 0.095 | 0.215 |
| FBA-1 | LH | clip_Xa | 5 | $0.0025 \pm 0.0018$ | 0.151 | 0.017 | 0.49 | 0.303 | 0.859 |
| FBA-1 | LH | clip_Xb | 5 | $0.0082 \pm 0.0025$ | 0.151 | 0.054 | 1.19 | 0.053 | 0.156 |
| FBA-1 | RH | bert_Xa | 6 | $0.0044 \pm 0.0029$ | 0.140 | 0.026 | 0.52 | 0.288 | 0.307 |
| FBA-1 | RH | bert_Xb | 6 | $0.0031 \pm 0.0030$ | 0.140 | 0.016 | 0.35 | 0.456 | 0.607 |
| FBA-1 | RH | gpt2_Xa | 6 | $0.0008 \pm 0.0019$ | 0.140 | 0.001 | 0.14 | 0.762 | 1.000 |
| FBA-1 | RH | gpt2_Xb | 6 | $0.0040 \pm 0.0027$ | 0.140 | 0.022 | 0.51 | 0.275 | 0.303 |
| FBA-1 | RH | clip_Xa | 6 | $-0.0006 \pm 0.0018$ | 0.140 | $-0.008$ | -0.11 | 0.810 | 1.000 |
| FBA-1 | RH | clip_Xb | 6 | $0.0021 \pm 0.0023$ | 0.140 | 0.010 | 0.32 | 0.474 | 0.498 |
| FBA-2 | LH | bert_Xa | 7 | $0.0115 \pm 0.0042$ | 0.180 | 0.051 | 0.91 | 0.056 | 0.086 |
| FBA-2 | LH | bert_Xb | 7 | $0.0125 \pm 0.0038$ | 0.180 | 0.060 | 1.08 | 0.034 | 0.086 |
| FBA-2 | LH | gpt2_Xa | 7 | $0.0077 \pm 0.0035$ | 0.180 | 0.032 | 0.72 | 0.126 | 0.184 |
| FBA-2 | LH | gpt2_Xb | 7 | $0.0163 \pm 0.0043$ | 0.180 | 0.084 | 1.23 | 0.021 | 0.043* |
| FBA-2 | LH | clip_Xa | 7 | $0.0038 \pm 0.0015$ | 0.180 | 0.020 | 0.81 | 0.074 | 0.344 |
| FBA-2 | LH | clip_Xb | 7 | $0.0119 \pm 0.0023$ | 0.180 | 0.069 | 1.68 | 0.007 | 0.054 |
| FBA-2 | RH | bert_Xa | 8 | $0.0169 \pm 0.0043$ | 0.238 | 0.066 | 1.23 | 0.015 | 0.027* |
| FBA-2 | RH | bert_Xb | 8 | $0.0157 \pm 0.0047$ | 0.238 | 0.060 | 1.05 | 0.028 | 0.061 |
| FBA-2 | RH | gpt2_Xa | 8 | $0.0130 \pm 0.0039$ | 0.238 | 0.048 | 1.06 | 0.026 | 0.054 |
| FBA-2 | RH | gpt2_Xb | 8 | $0.0216 \pm 0.0051$ | 0.238 | 0.082 | 1.33 | 0.011 | 0.027* |

*Continued on next page*

Table 2 – *continued from previous page*

| **ROI** | **H** | **Model** | **N** | $R^2_{\text{wiped}}$ **(mean ± SEM)** | $R^2_{\text{raw}}$ | **Ratio** | $g$ | $q_{\text{FDR}}$ | $q_{\text{SF}}$ |
|---|---|---|---|---|---|---|---|---|---|
| FBA-2 | RH | clip_Xa | 8 | $0.0042 \pm 0.0015$ | 0.238 | 0.018 | 0.91 | 0.039 | 0.344 |
| FBA-2 | RH | clip_Xb | 8 | $0.0120 \pm 0.0024$ | 0.238 | 0.049 | 1.56 | 0.006 | 0.031* |
| mTL-bodies | RH | bert_Xa | 2 | $-0.0035 \pm 0.0005$ | 0.050 | −0.073 | -0.00 | 0.147 | 1.000 |
| mTL-bodies | RH | bert_Xb | 2 | $-0.0030 \pm 0.0010$ | 0.050 | −0.059 | -0.00 | 0.307 | 1.000 |
| mTL-bodies | RH | gpt2_Xa | 2 | $-0.0023 \pm 0.0000$ | 0.050 | −0.052 | -0.00 | 0.007 | 1.000 |
| mTL-bodies | RH | gpt2_Xb | 2 | $0.0007 \pm 0.0002$ | 0.050 | 0.016 | 0.00 | 0.241 | 0.509 |
| mTL-bodies | RH | clip_Xa | 2 | $-0.0033 \pm 0.0028$ | 0.050 | −0.091 | -0.00 | 0.516 | 1.000 |
| mTL-bodies | RH | clip_Xb | 2 | $-0.0023 \pm 0.0006$ | 0.050 | −0.055 | -0.00 | 0.244 | 1.000 |
| *Face-selective* | | | | | | | | | |
| FFA-1 | LH | bert_Xa | 8 | $0.0080 \pm 0.0025$ | 0.168 | 0.042 | 0.99 | 0.033 | 0.027* |
| FFA-1 | LH | bert_Xb | 8 | $0.0057 \pm 0.0025$ | 0.168 | 0.028 | 0.73 | 0.091 | 0.086 |
| FFA-1 | LH | gpt2_Xa | 8 | $0.0028 \pm 0.0020$ | 0.168 | 0.008 | 0.42 | 0.309 | 0.537 |
| FFA-1 | LH | gpt2_Xb | 8 | $0.0071 \pm 0.0027$ | 0.168 | 0.034 | 0.83 | 0.057 | 0.072 |
| FFA-1 | LH | clip_Xa | 8 | $-0.0000 \pm 0.0014$ | 0.168 | −0.007 | -0.00 | 0.991 | 1.000 |
| FFA-1 | LH | clip_Xb | 8 | $0.0038 \pm 0.0019$ | 0.168 | 0.015 | 0.61 | 0.154 | 0.269 |
| FFA-1 | RH | bert_Xa | 8 | $0.0121 \pm 0.0025$ | 0.201 | 0.058 | 1.51 | 0.006 | 0.027* |
| FFA-1 | RH | bert_Xb | 8 | $0.0090 \pm 0.0025$ | 0.201 | 0.042 | 1.15 | 0.020 | 0.043* |
| FFA-1 | RH | gpt2_Xa | 8 | $0.0070 \pm 0.0020$ | 0.201 | 0.032 | 1.08 | 0.025 | 0.054 |
| FFA-1 | RH | gpt2_Xb | 8 | $0.0121 \pm 0.0027$ | 0.201 | 0.057 | 1.42 | 0.008 | 0.027* |
| FFA-1 | RH | clip_Xa | 8 | $0.0014 \pm 0.0011$ | 0.201 | 0.007 | 0.39 | 0.308 | 0.859 |
| FFA-1 | RH | clip_Xb | 8 | $0.0065 \pm 0.0011$ | 0.201 | 0.032 | 1.87 | 0.003 | 0.031* |
| FFA-2 | LH | bert_Xa | 7 | $0.0084 \pm 0.0071$ | 0.154 | −0.051 | 0.39 | 0.369 | 0.506 |
| FFA-2 | LH | bert_Xb | 7 | $0.0067 \pm 0.0066$ | 0.154 | −0.063 | 0.33 | 0.456 | 0.607 |
| FFA-2 | LH | gpt2_Xa | 7 | $0.0048 \pm 0.0049$ | 0.154 | −0.035 | 0.33 | 0.461 | 0.859 |
| FFA-2 | LH | gpt2_Xb | 7 | $0.0100 \pm 0.0061$ | 0.154 | −0.002 | 0.54 | 0.232 | 0.263 |

*Continued on next page*

Table 2 – *continued from previous page*

| **ROI** | **H** | **Model** | **N** | $R^2_{\text{wiped}}$ **(mean ± SEM)** | $R^2_{\text{raw}}$ | **Ratio** | $g$ | $q_{\text{FDR}}$ | $q_{\text{SF}}$ |
|---|---|---|---|---|---|---|---|---|---|
| FFA-2 | LH | clip_Xa | 7 | $-0.0005 \pm 0.0025$ | 0.154 | −0.064 | -0.06 | 0.872 | 1.000 |
| FFA-2 | LH | clip_Xb | 7 | $0.0042 \pm 0.0039$ | 0.154 | −0.039 | 0.36 | 0.408 | 0.498 |
| FFA-2 | RH | bert_Xa | 8 | $0.0077 \pm 0.0046$ | 0.168 | −0.022 | 0.52 | 0.216 | 0.281 |
| FFA-2 | RH | bert_Xb | 8 | $0.0063 \pm 0.0049$ | 0.168 | −0.037 | 0.41 | 0.344 | 0.562 |
| FFA-2 | RH | gpt2_Xa | 8 | $0.0059 \pm 0.0030$ | 0.168 | −0.004 | 0.61 | 0.158 | 0.322 |
| FFA-2 | RH | gpt2_Xb | 8 | $0.0116 \pm 0.0043$ | 0.168 | 0.032 | 0.84 | 0.056 | 0.043* |
| FFA-2 | RH | clip_Xa | 8 | $-0.0006 \pm 0.0021$ | 0.168 | −0.061 | -0.08 | 0.834 | 1.000 |
| FFA-2 | RH | clip_Xb | 8 | $0.0047 \pm 0.0034$ | 0.168 | −0.031 | 0.43 | 0.303 | 0.460 |
| OFA | LH | bert_Xa | 8 | $0.0001 \pm 0.0024$ | 0.120 | −0.016 | 0.01 | 0.981 | 0.911 |
| OFA | LH | bert_Xb | 8 | $-0.0013 \pm 0.0021$ | 0.120 | −0.027 | -0.20 | 0.629 | 1.000 |
| OFA | LH | gpt2_Xa | 8 | $-0.0021 \pm 0.0020$ | 0.120 | −0.035 | -0.33 | 0.427 | 1.000 |
| OFA | LH | gpt2_Xb | 8 | $0.0002 \pm 0.0024$ | 0.120 | −0.017 | 0.03 | 0.934 | 0.888 |
| OFA | LH | clip_Xa | 8 | $-0.0031 \pm 0.0011$ | 0.120 | −0.035 | -0.90 | 0.039 | 1.000 |
| OFA | LH | clip_Xb | 8 | $-0.0008 \pm 0.0014$ | 0.120 | −0.017 | -0.19 | 0.617 | 1.000 |
| OFA | RH | bert_Xa | 8 | $-0.0036 \pm 0.0028$ | 0.099 | −0.094 | -0.40 | 0.341 | 1.000 |
| OFA | RH | bert_Xb | 8 | $-0.0047 \pm 0.0027$ | 0.099 | −0.107 | -0.54 | 0.207 | 1.000 |
| OFA | RH | gpt2_Xa | 8 | $-0.0035 \pm 0.0017$ | 0.099 | −0.077 | -0.63 | 0.143 | 1.000 |
| OFA | RH | gpt2_Xb | 8 | $-0.0017 \pm 0.0023$ | 0.099 | −0.062 | -0.23 | 0.534 | 1.000 |
| OFA | RH | clip_Xa | 8 | $-0.0050 \pm 0.0013$ | 0.099 | −0.085 | -1.19 | 0.014 | 1.000 |
| OFA | RH | clip_Xb | 8 | $-0.0029 \pm 0.0017$ | 0.099 | −0.064 | -0.53 | 0.216 | 1.000 |
| *Place-selective* | | | | | | | | | |
| OPA | LH | bert_Xa | 8 | $0.0013 \pm 0.0015$ | 0.140 | 0.003 | 0.26 | 0.515 | 0.762 |
| OPA | LH | bert_Xb | 8 | $0.0013 \pm 0.0017$ | 0.140 | 0.002 | 0.25 | 0.568 | 0.908 |
| OPA | LH | gpt2_Xa | 8 | $-0.0015 \pm 0.0019$ | 0.140 | −0.022 | -0.25 | 0.536 | 1.000 |
| OPA | LH | gpt2_Xb | 8 | $0.0024 \pm 0.0023$ | 0.140 | 0.006 | 0.32 | 0.397 | 0.471 |

*Continued on next page*

Table 2 – *continued from previous page*

| **ROI** | **H** | **Model** | **N** | $R^2_{\text{wiped}}$ **(mean ± SEM)** | $R^2_{\text{raw}}$ | **Ratio** | $g$ | $q_{\text{FDR}}$ | $q_{\text{SF}}$ |
|---|---|---|---|---|---|---|---|---|---|
| OPA | LH | clip_Xa | 8 | $-0.0038 \pm 0.0008$ | 0.140 | $-0.030$ | -1.43 | 0.006 | 1.000 |
| OPA | LH | clip_Xb | 8 | $0.0012 \pm 0.0010$ | 0.140 | 0.007 | 0.38 | 0.364 | 0.493 |
| OPA | RH | bert_Xa | 8 | $0.0023 \pm 0.0018$ | 0.133 | 0.013 | 0.40 | 0.340 | 0.473 |
| OPA | RH | bert_Xb | 8 | $0.0023 \pm 0.0019$ | 0.133 | 0.011 | 0.38 | 0.366 | 0.602 |
| OPA | RH | gpt2_Xa | 8 | $-0.0013 \pm 0.0018$ | 0.133 | $-0.018$ | -0.22 | 0.576 | 1.000 |
| OPA | RH | gpt2_Xb | 8 | $0.0028 \pm 0.0024$ | 0.133 | 0.012 | 0.37 | 0.343 | 0.240 |
| OPA | RH | clip_Xa | 8 | $-0.0042 \pm 0.0013$ | 0.133 | $-0.034$ | -1.03 | 0.025 | 1.000 |
| OPA | RH | clip_Xb | 8 | $0.0011 \pm 0.0013$ | 0.133 | 0.006 | 0.26 | 0.499 | 0.557 |
| PPA | LH | bert_Xa | 8 | $0.0014 \pm 0.0026$ | 0.190 | 0.000 | 0.17 | 0.660 | 0.822 |
| PPA | LH | bert_Xb | 8 | $0.0001 \pm 0.0025$ | 0.190 | $-0.007$ | 0.02 | 0.955 | 0.959 |
| PPA | LH | gpt2_Xa | 8 | $-0.0006 \pm 0.0024$ | 0.190 | $-0.012$ | -0.07 | 0.855 | 1.000 |
| PPA | LH | gpt2_Xb | 8 | $0.0031 \pm 0.0030$ | 0.190 | 0.007 | 0.32 | 0.397 | 0.464 |
| PPA | LH | clip_Xa | 8 | $-0.0041 \pm 0.0014$ | 0.190 | $-0.026$ | -0.91 | 0.039 | 1.000 |
| PPA | LH | clip_Xb | 8 | $0.0004 \pm 0.0017$ | 0.190 | $-0.002$ | 0.07 | 0.842 | 0.760 |
| PPA | RH | bert_Xa | 8 | $0.0026 \pm 0.0022$ | 0.192 | 0.010 | 0.38 | 0.355 | 0.497 |
| PPA | RH | bert_Xb | 8 | $0.0010 \pm 0.0019$ | 0.192 | 0.002 | 0.16 | 0.683 | 0.959 |
| PPA | RH | gpt2_Xa | 8 | $-0.0004 \pm 0.0021$ | 0.192 | $-0.008$ | -0.07 | 0.857 | 1.000 |
| PPA | RH | gpt2_Xb | 8 | $0.0029 \pm 0.0024$ | 0.192 | 0.010 | 0.38 | 0.333 | 0.318 |
| PPA | RH | clip_Xa | 8 | $-0.0040 \pm 0.0014$ | 0.192 | $-0.021$ | -0.87 | 0.044 | 1.000 |
| PPA | RH | clip_Xb | 8 | $0.0005 \pm 0.0014$ | 0.192 | 0.002 | 0.11 | 0.754 | 0.760 |
| RSC | LH | bert_Xa | 8 | $0.0009 \pm 0.0038$ | 0.172 | $-0.020$ | 0.07 | 0.845 | 0.890 |
| RSC | LH | bert_Xb | 8 | $0.0004 \pm 0.0040$ | 0.172 | $-0.023$ | 0.03 | 0.951 | 0.959 |
| RSC | LH | gpt2_Xa | 8 | $-0.0002 \pm 0.0022$ | 0.172 | $-0.021$ | -0.03 | 0.929 | 1.000 |
| RSC | LH | gpt2_Xb | 8 | $0.0036 \pm 0.0029$ | 0.172 | 0.003 | 0.39 | 0.328 | 0.303 |
| RSC | LH | clip_Xa | 8 | $-0.0030 \pm 0.0021$ | 0.172 | $-0.034$ | -0.45 | 0.251 | 1.000 |
| RSC | LH | clip_Xb | 8 | $0.0010 \pm 0.0026$ | 0.172 | $-0.009$ | 0.12 | 0.754 | 0.760 |

*Continued on next page*

Table 2 – *continued from previous page*

| **ROI** | **H** | **Model** | **N** | $R^2_{\text{wiped}}$ **(mean ± SEM)** | $R^2_{\text{raw}}$ | **Ratio** | $g$ | $q_{\text{FDR}}$ | $q_{\text{SF}}$ |
|---|---|---|---|---|---|---|---|---|---|
| RSC | RH | bert_Xa | 8 | 0.0009 ± 0.0022 | 0.168 | 0.001 | 0.12 | 0.747 | 0.825 |
| RSC | RH | bert_Xb | 8 | 0.0005 ± 0.0024 | 0.168 | −0.002 | 0.06 | 0.895 | 0.959 |
| RSC | RH | gpt2_Xa | 8 | −0.0008 ± 0.0009 | 0.168 | −0.008 | -0.27 | 0.500 | 1.000 |
| RSC | RH | gpt2_Xb | 8 | 0.0030 ± 0.0011 | 0.168 | 0.016 | 0.81 | 0.058 | 0.072 |
| RSC | RH | clip_Xa | 8 | −0.0026 ± 0.0016 | 0.168 | −0.017 | -0.50 | 0.207 | 1.000 |
| RSC | RH | clip_Xb | 8 | 0.0016 ± 0.0019 | 0.168 | 0.009 | 0.27 | 0.492 | 0.557 |
| *Word-selective* | | | | | | | | | |
| OWFA | LH | bert_Xa | 8 | −0.0067 ± 0.0025 | 0.070 | −0.149 | -0.84 | 0.056 | 1.000 |
| OWFA | LH | bert_Xb | 8 | −0.0074 ± 0.0024 | 0.070 | −0.160 | -0.97 | 0.035 | 1.000 |
| OWFA | LH | gpt2_Xa | 8 | −0.0069 ± 0.0017 | 0.070 | −0.141 | -1.30 | 0.011 | 1.000 |
| OWFA | LH | gpt2_Xb | 8 | −0.0060 ± 0.0021 | 0.070 | −0.132 | -0.92 | 0.044 | 1.000 |
| OWFA | LH | clip_Xa | 8 | −0.0059 ± 0.0011 | 0.070 | −0.111 | -1.71 | 0.003 | 1.000 |
| OWFA | LH | clip_Xb | 8 | −0.0048 ± 0.0014 | 0.070 | −0.098 | -1.07 | 0.024 | 1.000 |
| OWFA | RH | bert_Xa | 8 | −0.0073 ± 0.0022 | 0.061 | −0.218 | -1.03 | 0.030 | 1.000 |
| OWFA | RH | bert_Xb | 8 | −0.0086 ± 0.0024 | 0.061 | −0.255 | -1.14 | 0.020 | 1.000 |
| OWFA | RH | gpt2_Xa | 8 | −0.0072 ± 0.0014 | 0.061 | −0.197 | -1.62 | 0.004 | 1.000 |
| OWFA | RH | gpt2_Xb | 8 | −0.0069 ± 0.0018 | 0.061 | −0.199 | -1.18 | 0.017 | 1.000 |
| OWFA | RH | clip_Xa | 8 | −0.0063 ± 0.0016 | 0.061 | −0.178 | -1.28 | 0.010 | 1.000 |
| OWFA | RH | clip_Xb | 8 | −0.0059 ± 0.0018 | 0.061 | −0.174 | -1.01 | 0.030 | 1.000 |
| VWFA-1 | LH | bert_Xa | 8 | 0.0008 ± 0.0016 | 0.122 | 0.001 | 0.17 | 0.660 | 0.762 |
| VWFA-1 | LH | bert_Xb | 8 | 0.0008 ± 0.0016 | 0.122 | 0.002 | 0.16 | 0.683 | 0.908 |
| VWFA-1 | LH | gpt2_Xa | 8 | −0.0016 ± 0.0013 | 0.122 | −0.026 | -0.39 | 0.358 | 1.000 |
| VWFA-1 | LH | gpt2_Xb | 8 | 0.0024 ± 0.0018 | 0.122 | 0.008 | 0.42 | 0.297 | 0.303 |
| VWFA-1 | LH | clip_Xa | 8 | −0.0024 ± 0.0017 | 0.122 | −0.028 | -0.45 | 0.251 | 1.000 |
| VWFA-1 | LH | clip_Xb | 8 | 0.0025 ± 0.0022 | 0.122 | 0.013 | 0.37 | 0.367 | 0.493 |

*Continued on next page*

Table 2 – *continued from previous page*

| **ROI** | **H** | **Model** | **N** | $R^2_{\text{wiped}}$ **(mean ± SEM)** | $R^2_{\text{raw}}$ | **Ratio** | $g$ | $q_{\text{FDR}}$ | $q_{\text{SF}}$ |
|---|---|---|---|---|---|---|---|---|---|
| VWFA-1 | RH | bert_Xa | 8 | $0.0062 \pm 0.0016$ | 0.156 | 0.041 | 1.18 | 0.017 | 0.048* |
| VWFA-1 | RH | bert_Xb | 8 | $0.0056 \pm 0.0018$ | 0.156 | 0.037 | 0.98 | 0.034 | 0.081 |
| VWFA-1 | RH | gpt2_Xa | 8 | $0.0025 \pm 0.0017$ | 0.156 | 0.009 | 0.44 | 0.297 | 0.537 |
| VWFA-1 | RH | gpt2_Xb | 8 | $0.0070 \pm 0.0023$ | 0.156 | 0.040 | 0.98 | 0.036 | 0.027* |
| VWFA-1 | RH | clip_Xa | 8 | $-0.0009 \pm 0.0009$ | 0.156 | $-0.006$ | -0.31 | 0.418 | 1.000 |
| VWFA-1 | RH | clip_Xb | 8 | $0.0043 \pm 0.0012$ | 0.156 | 0.025 | 1.08 | 0.024 | 0.031* |
| VWFA-2 | LH | bert_Xa | 8 | $0.0043 \pm 0.0026$ | 0.114 | 0.014 | 0.53 | 0.210 | 0.281 |
| VWFA-2 | LH | bert_Xb | 8 | $0.0071 \pm 0.0034$ | 0.114 | 0.035 | 0.66 | 0.120 | 0.215 |
| VWFA-2 | LH | gpt2_Xa | 8 | $0.0012 \pm 0.0018$ | 0.114 | $-0.013$ | 0.21 | 0.593 | 1.000 |
| VWFA-2 | LH | gpt2_Xb | 8 | $0.0076 \pm 0.0028$ | 0.114 | 0.046 | 0.85 | 0.056 | 0.083 |
| VWFA-2 | LH | clip_Xa | 8 | $-0.0010 \pm 0.0017$ | 0.114 | $-0.025$ | -0.18 | 0.636 | 1.000 |
| VWFA-2 | LH | clip_Xb | 8 | $0.0071 \pm 0.0033$ | 0.114 | 0.041 | 0.68 | 0.113 | 0.119 |
| VWFA-2 | RH | bert_Xa | 5 | $0.0015 \pm 0.0015$ | 0.141 | 0.011 | 0.36 | 0.454 | 0.602 |
| VWFA-2 | RH | bert_Xb | 5 | $0.0011 \pm 0.0015$ | 0.141 | 0.009 | 0.27 | 0.584 | 0.724 |
| VWFA-2 | RH | gpt2_Xa | 5 | $0.0008 \pm 0.0013$ | 0.141 | 0.007 | 0.22 | 0.620 | 1.000 |
| VWFA-2 | RH | gpt2_Xb | 5 | $0.0054 \pm 0.0016$ | 0.141 | 0.040 | 1.18 | 0.056 | 0.123 |
| VWFA-2 | RH | clip_Xa | 5 | $-0.0014 \pm 0.0019$ | 0.141 | $-0.008$ | -0.26 | 0.560 | 1.000 |
| VWFA-2 | RH | clip_Xb | 5 | $0.0033 \pm 0.0017$ | 0.141 | 0.026 | 0.69 | 0.203 | 0.397 |
| mfs-words | LH | bert_Xa | 8 | $0.0026 \pm 0.0040$ | 0.127 | $-0.007$ | 0.20 | 0.615 | 0.828 |
| mfs-words | LH | bert_Xb | 8 | $0.0010 \pm 0.0039$ | 0.127 | $-0.022$ | 0.08 | 0.861 | 0.959 |
| mfs-words | LH | gpt2_Xa | 8 | $-0.0027 \pm 0.0024$ | 0.127 | $-0.050$ | -0.36 | 0.389 | 1.000 |
| mfs-words | LH | gpt2_Xb | 8 | $0.0006 \pm 0.0032$ | 0.127 | $-0.024$ | 0.06 | 0.879 | 0.852 |
| mfs-words | LH | clip_Xa | 8 | $-0.0014 \pm 0.0021$ | 0.127 | $-0.026$ | -0.22 | 0.560 | 1.000 |
| mfs-words | LH | clip_Xb | 8 | $0.0013 \pm 0.0028$ | 0.127 | $-0.008$ | 0.15 | 0.705 | 0.859 |
| mfs-words | RH | bert_Xa | 6 | $0.0022 \pm 0.0030$ | 0.122 | 0.005 | 0.26 | 0.575 | 0.824 |
| mfs-words | RH | bert_Xb | 6 | $0.0003 \pm 0.0030$ | 0.122 | $-0.012$ | 0.03 | 0.951 | 0.959 |

*Continued on next page*

Table 2 – *continued from previous page*

| ROI | H | Model | N | $R^2_{\text{wiped}}$ (mean ± SEM) | $R^2_{\text{raw}}$ | Ratio | $g$ | $q_{\text{FDR}}$ | $q_{\text{SF}}$ |
|---|---|---|---|---|---|---|---|---|---|
| mfs-words | RH | gpt2_Xa | 6 | $-0.0020 \pm 0.0021$ | 0.122 | −0.030 | -0.33 | 0.479 | 1.000 |
| mfs-words | RH | gpt2_Xb | 6 | $0.0006 \pm 0.0025$ | 0.122 | −0.009 | 0.09 | 0.843 | 0.852 |
| mfs-words | RH | clip_Xa | 6 | $-0.0019 \pm 0.0017$ | 0.122 | −0.025 | -0.40 | 0.361 | 1.000 |
| mfs-words | RH | clip_Xb | 6 | $0.0004 \pm 0.0023$ | 0.122 | −0.009 | 0.05 | 0.880 | 0.921 |
| *Early visual (retinotopic)* | | | | | | | | | |
| V1d | LH | bert_Xa | 8 | $-0.0100 \pm 0.0015$ | 0.065 | −0.175 | -2.15 | 0.001 | 1.000 |
| V1d | LH | bert_Xb | 8 | $-0.0097 \pm 0.0013$ | 0.065 | −0.168 | -2.30 | <0.001 | 1.000 |
| V1d | LH | gpt2_Xa | 8 | $-0.0098 \pm 0.0012$ | 0.065 | −0.167 | -2.51 | <0.001 | 1.000 |
| V1d | LH | gpt2_Xb | 8 | $-0.0089 \pm 0.0012$ | 0.065 | −0.151 | -2.28 | 0.001 | 1.000 |
| V1d | LH | clip_Xa | 8 | $-0.0069 \pm 0.0010$ | 0.065 | −0.117 | -2.07 | 0.001 | 1.000 |
| V1d | LH | clip_Xb | 8 | $-0.0063 \pm 0.0009$ | 0.065 | −0.110 | -2.25 | 0.001 | 1.000 |
| V1d | RH | bert_Xa | 8 | $-0.0086 \pm 0.0014$ | 0.064 | −0.157 | -1.92 | 0.002 | 1.000 |
| V1d | RH | bert_Xb | 8 | $-0.0084 \pm 0.0015$ | 0.064 | −0.153 | -1.74 | 0.003 | 1.000 |
| V1d | RH | gpt2_Xa | 8 | $-0.0081 \pm 0.0011$ | 0.064 | −0.143 | -2.29 | <0.001 | 1.000 |
| V1d | RH | gpt2_Xb | 8 | $-0.0076 \pm 0.0012$ | 0.064 | −0.136 | -1.94 | 0.002 | 1.000 |
| V1d | RH | clip_Xa | 8 | $-0.0072 \pm 0.0011$ | 0.064 | −0.125 | -2.12 | 0.001 | 1.000 |
| V1d | RH | clip_Xb | 8 | $-0.0069 \pm 0.0007$ | 0.064 | −0.120 | -3.07 | <0.001 | 1.000 |
| V1v | LH | bert_Xa | 8 | $-0.0102 \pm 0.0008$ | 0.060 | −0.183 | -3.91 | <0.001 | 1.000 |
| V1v | LH | bert_Xb | 8 | $-0.0101 \pm 0.0008$ | 0.060 | −0.180 | -3.82 | <0.001 | 1.000 |
| V1v | LH | gpt2_Xa | 8 | $-0.0097 \pm 0.0007$ | 0.060 | −0.170 | -4.25 | <0.001 | 1.000 |
| V1v | LH | gpt2_Xb | 8 | $-0.0092 \pm 0.0007$ | 0.060 | −0.162 | -4.31 | <0.001 | 1.000 |
| V1v | LH | clip_Xa | 8 | $-0.0069 \pm 0.0008$ | 0.060 | −0.121 | -2.69 | <0.001 | 1.000 |
| V1v | LH | clip_Xb | 8 | $-0.0066 \pm 0.0008$ | 0.060 | −0.116 | -2.69 | <0.001 | 1.000 |
| V1v | RH | bert_Xa | 8 | $-0.0090 \pm 0.0015$ | 0.063 | −0.180 | -1.94 | 0.002 | 1.000 |
| V1v | RH | bert_Xb | 8 | $-0.0090 \pm 0.0015$ | 0.063 | −0.181 | -1.91 | 0.002 | 1.000 |

*Continued on next page*

Table 2 – *continued from previous page*

| ROI | H | Model | N | $R^2_{\text{wiped}}$ (mean ± SEM) | $R^2_{\text{raw}}$ | Ratio | $g$ | $q_{\text{FDR}}$ | $q_{\text{SF}}$ |
|---|---|---|---|---|---|---|---|---|---|
| V1v | RH | gpt2_Xa | 8 | $-0.0076 \pm 0.0010$ | 0.063 | $-0.147$ | -2.49 | <0.001 | 1.000 |
| V1v | RH | gpt2_Xb | 8 | $-0.0068 \pm 0.0011$ | 0.063 | $-0.136$ | -2.02 | 0.002 | 1.000 |
| V1v | RH | clip_Xa | 8 | $-0.0062 \pm 0.0008$ | 0.063 | $-0.115$ | -2.55 | <0.001 | 1.000 |
| V1v | RH | clip_Xb | 8 | $-0.0055 \pm 0.0008$ | 0.063 | $-0.107$ | -2.26 | 0.001 | 1.000 |
| V2d | LH | bert_Xa | 8 | $-0.0107 \pm 0.0013$ | 0.048 | $-0.251$ | -2.57 | <0.001 | 1.000 |
| V2d | LH | bert_Xb | 8 | $-0.0103 \pm 0.0012$ | 0.048 | $-0.243$ | -2.79 | <0.001 | 1.000 |
| V2d | LH | gpt2_Xa | 8 | $-0.0094 \pm 0.0010$ | 0.048 | $-0.215$ | -3.00 | <0.001 | 1.000 |
| V2d | LH | gpt2_Xb | 8 | $-0.0083 \pm 0.0010$ | 0.048 | $-0.192$ | -2.63 | <0.001 | 1.000 |
| V2d | LH | clip_Xa | 8 | $-0.0080 \pm 0.0010$ | 0.048 | $-0.184$ | -2.54 | <0.001 | 1.000 |
| V2d | LH | clip_Xb | 8 | $-0.0076 \pm 0.0009$ | 0.048 | $-0.174$ | -2.77 | <0.001 | 1.000 |
| V2d | RH | bert_Xa | 8 | $-0.0094 \pm 0.0008$ | 0.051 | $-0.204$ | -3.90 | <0.001 | 1.000 |
| V2d | RH | bert_Xb | 8 | $-0.0097 \pm 0.0008$ | 0.051 | $-0.208$ | -3.72 | <0.001 | 1.000 |
| V2d | RH | gpt2_Xa | 8 | $-0.0097 \pm 0.0007$ | 0.051 | $-0.201$ | -4.12 | <0.001 | 1.000 |
| V2d | RH | gpt2_Xb | 8 | $-0.0092 \pm 0.0008$ | 0.051 | $-0.193$ | -3.72 | <0.001 | 1.000 |
| V2d | RH | clip_Xa | 8 | $-0.0086 \pm 0.0011$ | 0.051 | $-0.184$ | -2.45 | <0.001 | 1.000 |
| V2d | RH | clip_Xb | 8 | $-0.0084 \pm 0.0009$ | 0.051 | $-0.179$ | -3.05 | <0.001 | 1.000 |
| V2v | LH | bert_Xa | 8 | $-0.0078 \pm 0.0010$ | 0.067 | $-0.135$ | -2.37 | <0.001 | 1.000 |
| V2v | LH | bert_Xb | 8 | $-0.0075 \pm 0.0009$ | 0.067 | $-0.128$ | -2.50 | <0.001 | 1.000 |
| V2v | LH | gpt2_Xa | 8 | $-0.0080 \pm 0.0010$ | 0.067 | $-0.137$ | -2.50 | <0.001 | 1.000 |
| V2v | LH | gpt2_Xb | 8 | $-0.0072 \pm 0.0010$ | 0.067 | $-0.124$ | -2.19 | 0.001 | 1.000 |
| V2v | LH | clip_Xa | 8 | $-0.0066 \pm 0.0009$ | 0.067 | $-0.110$ | -2.33 | <0.001 | 1.000 |
| V2v | LH | clip_Xb | 8 | $-0.0057 \pm 0.0010$ | 0.067 | $-0.098$ | -1.76 | 0.003 | 1.000 |
| V2v | RH | bert_Xa | 8 | $-0.0084 \pm 0.0016$ | 0.060 | $-0.180$ | -1.65 | 0.004 | 1.000 |
| V2v | RH | bert_Xb | 8 | $-0.0084 \pm 0.0017$ | 0.060 | $-0.177$ | -1.55 | 0.005 | 1.000 |
| V2v | RH | gpt2_Xa | 8 | $-0.0070 \pm 0.0011$ | 0.060 | $-0.148$ | -2.08 | 0.001 | 1.000 |
| V2v | RH | gpt2_Xb | 8 | $-0.0062 \pm 0.0012$ | 0.060 | $-0.133$ | -1.62 | 0.005 | 1.000 |

*Continued on next page*

Table 2 – *continued from previous page*

| **ROI** | **H** | **Model** | **N** | $R^2_{\text{wiped}}$ **(mean ± SEM)** | $R^2_{\text{raw}}$ | **Ratio** | $g$ | $q_{\text{FDR}}$ | $q_{\text{SF}}$ |
|---|---|---|---|---|---|---|---|---|---|
| V2v | RH | clip_Xa | 8 | $-0.0060 \pm 0.0010$ | 0.060 | $-0.126$ | -1.99 | 0.002 | 1.000 |
| V2v | RH | clip_Xb | 8 | $-0.0054 \pm 0.0011$ | 0.060 | $-0.116$ | -1.54 | 0.006 | 1.000 |
| V3d | LH | bert_Xa | 8 | $-0.0091 \pm 0.0012$ | 0.050 | $-0.212$ | -2.38 | <0.001 | 1.000 |
| V3d | LH | bert_Xb | 8 | $-0.0090 \pm 0.0011$ | 0.050 | $-0.210$ | -2.51 | <0.001 | 1.000 |
| V3d | LH | gpt2_Xa | 8 | $-0.0088 \pm 0.0010$ | 0.050 | $-0.202$ | -2.83 | <0.001 | 1.000 |
| V3d | LH | gpt2_Xb | 8 | $-0.0076 \pm 0.0012$ | 0.050 | $-0.180$ | -1.95 | 0.002 | 1.000 |
| V3d | LH | clip_Xa | 8 | $-0.0076 \pm 0.0009$ | 0.050 | $-0.172$ | -2.66 | <0.001 | 1.000 |
| V3d | LH | clip_Xb | 8 | $-0.0070 \pm 0.0009$ | 0.050 | $-0.162$ | -2.44 | <0.001 | 1.000 |
| V3d | RH | bert_Xa | 8 | $-0.0090 \pm 0.0008$ | 0.052 | $-0.187$ | -3.50 | <0.001 | 1.000 |
| V3d | RH | bert_Xb | 8 | $-0.0092 \pm 0.0008$ | 0.052 | $-0.192$ | -3.52 | <0.001 | 1.000 |
| V3d | RH | gpt2_Xa | 8 | $-0.0079 \pm 0.0005$ | 0.052 | $-0.158$ | -4.73 | <0.001 | 1.000 |
| V3d | RH | gpt2_Xb | 8 | $-0.0076 \pm 0.0005$ | 0.052 | $-0.154$ | -4.99 | <0.001 | 1.000 |
| V3d | RH | clip_Xa | 8 | $-0.0082 \pm 0.0009$ | 0.052 | $-0.165$ | -3.01 | <0.001 | 1.000 |
| V3d | RH | clip_Xb | 8 | $-0.0077 \pm 0.0006$ | 0.052 | $-0.158$ | -3.87 | <0.001 | 1.000 |
| V3v | LH | bert_Xa | 8 | $-0.0070 \pm 0.0011$ | 0.061 | $-0.138$ | -2.05 | 0.002 | 1.000 |
| V3v | LH | bert_Xb | 8 | $-0.0066 \pm 0.0010$ | 0.061 | $-0.130$ | -2.06 | 0.002 | 1.000 |
| V3v | LH | gpt2_Xa | 8 | $-0.0074 \pm 0.0012$ | 0.061 | $-0.141$ | -1.99 | 0.001 | 1.000 |
| V3v | LH | gpt2_Xb | 8 | $-0.0061 \pm 0.0014$ | 0.061 | $-0.121$ | -1.42 | 0.008 | 1.000 |
| V3v | LH | clip_Xa | 8 | $-0.0058 \pm 0.0011$ | 0.061 | $-0.114$ | -1.72 | 0.003 | 1.000 |
| V3v | LH | clip_Xb | 8 | $-0.0045 \pm 0.0013$ | 0.061 | $-0.096$ | -1.11 | 0.023 | 1.000 |
| V3v | RH | bert_Xa | 8 | $-0.0086 \pm 0.0014$ | 0.057 | $-0.163$ | -1.89 | 0.002 | 1.000 |
| V3v | RH | bert_Xb | 8 | $-0.0087 \pm 0.0016$ | 0.057 | $-0.167$ | -1.74 | 0.003 | 1.000 |
| V3v | RH | gpt2_Xa | 8 | $-0.0071 \pm 0.0011$ | 0.057 | $-0.136$ | -2.05 | 0.001 | 1.000 |
| V3v | RH | gpt2_Xb | 8 | $-0.0062 \pm 0.0014$ | 0.057 | $-0.121$ | -1.42 | 0.008 | 1.000 |
| V3v | RH | clip_Xa | 8 | $-0.0063 \pm 0.0010$ | 0.057 | $-0.118$ | -2.07 | 0.001 | 1.000 |
| V3v | RH | clip_Xb | 8 | $-0.0058 \pm 0.0014$ | 0.057 | $-0.112$ | -1.30 | 0.011 | 1.000 |

*Continued on next page*

Table 2 – *continued from previous page*

| **ROI** | **H** | **Model** | **N** | $R^2_{\text{wiped}}$ **(mean ± SEM)** | $R^2_{\text{raw}}$ | **Ratio** | $g$ | $q_{\text{FDR}}$ | $q_{\text{SF}}$ |
|---|---|---|---|---|---|---|---|---|---|
| hV4 | LH | bert_Xa | 8 | $-0.0054 \pm 0.0008$ | 0.079 | $-0.079$ | -2.16 | 0.001 | 1.000 |
| hV4 | LH | bert_Xb | 8 | $-0.0060 \pm 0.0008$ | 0.079 | $-0.086$ | -2.46 | <0.001 | 1.000 |
| hV4 | LH | gpt2_Xa | 8 | $-0.0062 \pm 0.0010$ | 0.079 | $-0.088$ | -1.96 | 0.001 | 1.000 |
| hV4 | LH | gpt2_Xb | 8 | $-0.0051 \pm 0.0012$ | 0.079 | $-0.073$ | -1.30 | 0.011 | 1.000 |
| hV4 | LH | clip_Xa | 8 | $-0.0054 \pm 0.0009$ | 0.079 | $-0.078$ | -1.80 | 0.002 | 1.000 |
| hV4 | LH | clip_Xb | 8 | $-0.0041 \pm 0.0012$ | 0.079 | $-0.061$ | -1.10 | 0.023 | 1.000 |
| hV4 | RH | bert_Xa | 8 | $-0.0064 \pm 0.0011$ | 0.070 | $-0.107$ | -1.75 | 0.003 | 1.000 |
| hV4 | RH | bert_Xb | 8 | $-0.0072 \pm 0.0012$ | 0.070 | $-0.120$ | -1.85 | 0.003 | 1.000 |
| hV4 | RH | gpt2_Xa | 8 | $-0.0063 \pm 0.0008$ | 0.070 | $-0.101$ | -2.62 | <0.001 | 1.000 |
| hV4 | RH | gpt2_Xb | 8 | $-0.0053 \pm 0.0009$ | 0.070 | $-0.087$ | -1.76 | 0.003 | 1.000 |
| hV4 | RH | clip_Xa | 8 | $-0.0058 \pm 0.0011$ | 0.070 | $-0.095$ | -1.62 | 0.003 | 1.000 |
| hV4 | RH | clip_Xb | 8 | $-0.0051 \pm 0.0011$ | 0.070 | $-0.084$ | -1.50 | 0.007 | 1.000 |

H = hemisphere; Model = language/vision model and variant (Xa or Xb); N = number of subjects.
* significant by sign-flip permutation test after BH-FDR correction ($q_{\text{SF}} < 0.05$).
$q_{\text{FDR}}$: BH-corrected $p$-value from one-sample $t$-test across all ROI × hemisphere × model tests.
Ratio = $R^2_{\text{wiped}}/R^2_{\text{raw}}$. $g$ = Hedges' $g$ (effect size).
